# Rewiring of the phosphoproteome executes two meiotic divisions

**DOI:** 10.1101/2023.09.22.559006

**Authors:** Lori B. Koch, Christos Spanos, Van Kelly, Tony Ly, Adele L. Marston

## Abstract

The cell cycle is ordered by a controlled network of kinases and phosphatases. To generate gametes via meiosis, two distinct and sequential chromosome segregation events occur without an intervening S phase. How canonical cell cycle controls are modified for meiosis is not well understood. Here, using highly synchronous budding yeast populations, we reveal how the global proteome and phosphoproteome changes during the meiotic divisions. While protein abundance changes are limited to key cell cycle regulators, dynamic phosphorylation changes are pervasive. Our data indicate that two waves of cyclin-dependent kinase and Polo (Cdc5^Polo^) kinase activity drive successive meiotic divisions. These two distinct waves of phosphorylation are ensured by the meiosis-specific Spo13^Meikin^ protein, which rewires the phosphoproteome. Spo13^Meikin^ binds to Cdc5^Polo^ to promote phosphorylation of a subset of substrates in meiosis I containing a newly identified motif, which we define as the Spo13^Meikin^ -Cdc5^Polo^ consensus phosphorylation motif. Overall, our findings reveal that a master regulator of meiosis redirects the activity of a kinase to change the phosphorylation landscape and elicit a developmental cascade.

## Introduction

In mitotically dividing cells, ploidy is maintained by strict alternation of DNA replication in synthesis (S) phase and chromosome segregation in mitosis (M phase). The oscillating nature of the cell cycle is achieved through an inter-dependent combination of irreversible proteolytic degradation and reversible post-translational modifications. The major drivers of the cell cycle are cyclin-dependent kinases (Cdks). Cdk activity is low in G1, rises through S phase and peaks in mitosis after which Cdks are inactivated, triggering exit from mitosis and return to the G1 state. This state of low Cdk activity in G1 allows for the resetting of replication origins and entry into the next S phase. The events of mitotic exit are ordered by sequential inactivation of kinases, activation of phosphatases and selective dephosphorylation of particular substrates. In mammalian cells, proteolysis of cyclin B initiates mitotic exit, and thereafter the order of dephosphorylation is established by PP1 and PP2A phosphatase selectively dephosphorylating threonine over serine-centred sites (Holder *et al*, 2020; Kruse *et al*, 2020). In budding yeast, mitotic exit is ordered by sequential dephosphorylation of Cdk and Polo kinase phosphorylation sites and by the substrate preference of the phosphatase Cdc14 (Touati *et al*, 2018, 2019; Visintin *et al*, 1998).

The maintenance of ploidy in mitosis contrasts with meiosis where the goal is to generate gametes with half the ploidy of the parental cell. This is achieved by two consecutive chromosome segregation events (M phases), meiosis I and meiosis II, which follow a single S phase. Furthermore, meiosis I is a unique type of segregation event because homologs, rather than sister chromatids, are separated. This raises two key questions. First, how is the cell cycle machinery remodelled to direct the unique pattern of segregation in meiosis I, and second, what ensures that meiosis I is followed by another M phase, meiosis II, rather than S phase as in the canonical cell cycle?

The meiotic divisions are distinguished from mitotic division by three key features. First sister kinetochores attach to microtubules from the same, rather than opposite, poles in meiosis I (mono-orientation). Second, sister chromatid cohesion is lost in two steps: from chromosome arms in meiosis I and from centromeres only in meiosis II. Third, there are two successive chromosome segregation events (two divisions). The Meikin family of meiosis-specific regulators (Spo13 in budding yeast, Moa1 in fission yeast, Matrimony in *Drosophila* and Meikin in mammals) are master regulators which establish these key events of meiosis (Galander & Marston, 2020). Although Meikins are poorly conserved at the sequence level, all four proteins associate with Polo kinase through its polo binding domain (PBD) and, where tested, this interaction is required for their meiotic functions (Matos *et al*, 2008; Galander *et al*, 2019; Kim *et al*, 2015; Bonner *et al*, 2020). This suggests that Meikins define the meiotic programme by influencing Polo-dependent phosphorylation. Indeed, Polo plays several essential functions in meiosis, though only some of them overlap with Meikin. For example, in budding yeast, depletion of Cdc5^Polo^ causes a loss of monoorientation and arrest prior to meiotic division, while *spo13Δ* cells also lose monoorientation but undergo a meiotic division (Lee & Amon, 2003; Lee *et al*, 2004; Clyne *et al*, 2003; Katis *et al*, 2004). Spo13^Meikin^ also influences the activity of other kinases to prevent meiosis II events, such as loss of centromeric cohesion and spore formation, occurring prematurely, though it is unclear if this is direct or through its association with Cdc5^Polo^ (Oz *et al*, 2022; Galander *et al*, 2019).

In addition to meiosis-specific Polo regulation, the oscillations of Cdk activity are also expected to change to avoid completely exiting M phase after the first meiotic division. Indeed, early studies in the frog *Xenopus laevis* indicated that Cdk activity declines after anaphase I, but not completely, and that it quickly rises again during the transition to metaphase II (Furuno *et al*, 1994; Iwabuchi *et al*, 2000). More recently, analysis of sea star oocytes provided evidence that a rise in PP2A-B55 phosphatase activity at the meiosis I to II transition leads to the selective dephosphorylation of threonine-centered Cdk phosphorylation sites after meiosis I (Swartz *et al*, 2021). In budding yeast, progression through the meiotic divisions requires two major kinases, Cdc28^Cdk^ and a Cdk-related kinase Ime2 (Benjamin *et al*, 2003; Schindler & Winter, 2006). Distinct combinations of Cdk-cyclin complexes are active in each of the two meiotic divisions (Carlile & Amon, 2008). Both Cdk and Ime2 are required to prevent loading of the replicative helicase at the meiosis I to II transition (Phizicky *et al*, 2018; Holt *et al*, 2007). Further evidence for specialized control of the meiosis I to meiosis II transition came from studies of the budding yeast Cdc14 phosphatase which is critical for Cdk inactivation at mitotic exit (Visintin *et al*, 1998), but not at the meiosis I to II transition where it plays a specific role in licensing a second round of spindle pole body duplication (Marston *et al*, 2003; Bizzari & Marston, 2011; Fox *et al*, 2017). Together, these observations suggest that Cdk inactivation and phosphatase activation do occur at the meiosis I-meiosis II transition but only partially, achieving a kinase:phosphatase balance that maintains a subset of phosphorylations important for a subsequent M phase, rather than S phase, but how this is achieved is unclear.

To begin to understand the signalling events that create the meiotic programme and allow two sequential meiotic divisions, we have characterized the proteome and phosphoproteome of budding yeast cells synchronously undergoing the meiotic divisions. A combination of hierarchical clustering and motif analysis allowed us to infer the timing and activity of key meiotic kinases with implications for the control of meiosis. By generating a second, matched time-resolved dataset of *spo13Δ* cells, we defined how a master regulator establishes the meiotic programme. Finally, by comparing the proteome and phosphoproteome of metaphase I-arrested wild type, *spo13Δ* and *spo13-m2* cells, which abolish Cdc5^Polo^ binding, we reveal a motif and substrates targeted by Spo13^Meikin^-Cdc5^Polo^ in meiosis I.

## Results

### A high-resolution proteome and phosphoproteome of the meiotic divisions

To discover the proteins and phosphorylation events at each stage of the meiotic divisions in budding yeast, we released cells from a prophase I block (Carlile & Amon, 2008) and confirmed synchrony by spindle morphology (Fig 1A and Appendix Fig S1A). The total proteome and phosphoproteome was analysed by tandem mass tag (TMT) mass spectrometry at 15 minute intervals spanning the meiotic divisions and prophase “time zero” (Fig 1A; see Methods). In each of two biological replicates, we identified ∼3,600-4,000 proteins with 3,499 overlap between replicates (Fig 1B). From singly phosphorylated peptides detected in at least two sequential timepoints we quantified ∼6,700-8,700 phospho-sites in each timecourse, with 4,551 sites detected in both replicates (Appendix Fig S1B). To more accurately analyse the dynamics of phosphorylation, we normalised the phospho-site abundances to their corresponding total protein abundance and quantified ∼5,500-7,100 high confidence sites (localization score >0.75; (Olsen *et al*, 2006)) per replicate, of which 3,877 were found in both replicates (Fig 1B and Appendix Fig S1B). All subsequent analyses were restricted to the 3,499 proteins and 3,877 phospho-sites detected in both replicates, with phospho-sites normalized to their protein abundance (see Methods for details of normalization and batch correction procedures used).

**Figure 1.**
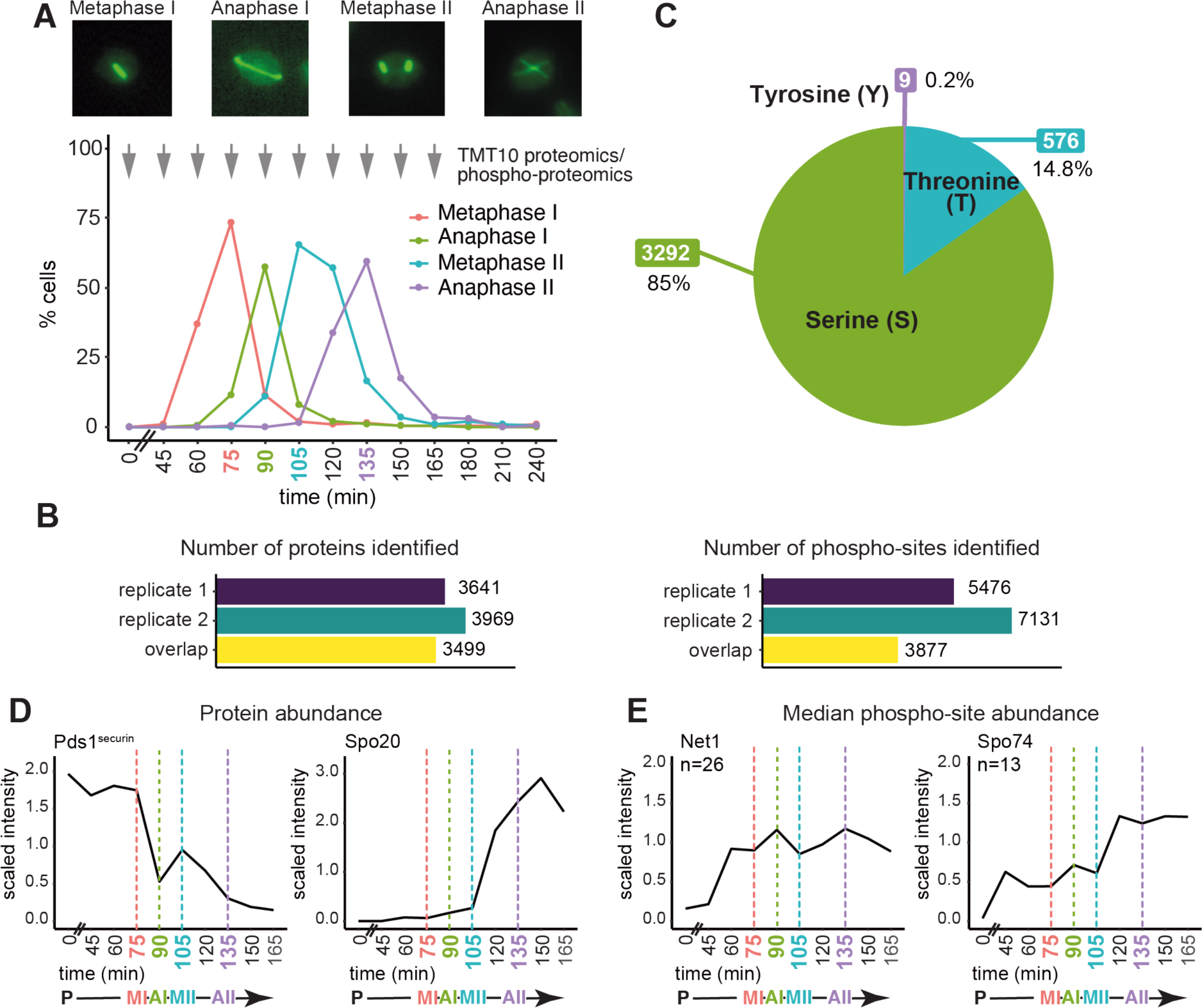
Phospho-proteomics of a synchronous meiotic division cycle. A. Example images and quantification of spindle immunofluorescence at the indicated timepoints after release of wild-type cells from a prophase I block. (n=200 cells per timepoint). Arrows indicate time of harvesting for TMT10 proteomics and phospho-proteomics. B. Total number of proteins (left) and phospho-sites (right) quantified for each of two biological replicate wild type TMT10 time-courses and those common to both time-courses (overlap). C. The proportion of phospho-sites centred on serine, threonine or tyrosine. D. Median protein abundance across the timecourse for selected proteins Pds1^securin^ and Spo20. E. Median phospho-site abundance for all detected phospho-sites for example proteins Net1 and Spo74.

The numbers of protein and phospho-sites detected, and their distributions, were comparable at all timepoints (Appendix Fig S1C and D). Slightly fewer phospho-sites were detected in prophase, indicating *de novo* phosphorylation thereafter (Appendix Fig S1D). Serine-centred phospho-sites (85%) comprised the majority of the meiotic phosphoproteome with threonine (15%) and tyrosine (<1%) less abundant, comparable to mitosis (Touati *et al*, 2018) (Fig 1C). We confirmed that our dataset has sufficient resolution to capture the expected dynamics of proteins and their phosphorylation over the two meiotic divisions. Two waves of securin (Pds1) destruction were detected (in anaphase I and II, (Salah & Nasmyth, 2000) and the Spo20 prospore membrane formation protein increased in abundance specifically during meiosis II (Neiman, 1998) (Fig 1D). Similarly, phosphorylation of the nucleolar protein Net1 peaked in both anaphase I and anaphase II, when its inhibitory association with the cell cycle regulatory phosphatase Cdc14 is relieved (Marston *et al*, 2003; Buonomo *et al*, 2003), while phosphorylation of the meiosis-specific centrosome protein Spo74 peaked during meiosis II, when it carries out its role in prospore membrane formation (Nickas *et al*, 2003; Ubersax *et al*, 2003) (Fig 1E). Together, these analyses confirm acquisition of a high-quality proteome and phosphoproteome of the meiotic divisions.

### Modest changes in global protein dynamics across the meiotic divisions

To identify the most dynamic meiotic proteins, we compared each of the 9 timepoints spanning the divisions with prophase time zero. A median fold change in abundance greater than 1.5 and a t-test value of p< 0.05 in at least one comparison defined 297 proteins as significantly dynamic, representing ∼8% of the meiotic proteome (Fig EV1A). To reveal their trends in abundance across the time-course in an unbiased manner, we used hierarchical clustering (Fig 2A and B). This grouped the dynamic proteins into six clusters, indicating enrichment of distinct subsets of proteins at all stages of meiosis. A group of proteins enriched specifically at prophase (cluster 3) was associated with GO terms involved in metabolism (Fig 2C). Cluster 2 proteins were abundant from prophase until anaphase I and involved in chromosome pairing, cohesion, and recombination. Cluster 5 proteins were abundant in meiosis I and II and associated with meiotic cell cycle and sporulation GO terms. Clusters 1 and 4 had high abundance in meiosis II and included proteins involved in sister-chromatid cohesion/anaphase-promoting complex (APC) activity and spore wall assembly (Fig 2C).

**Figure 2.**
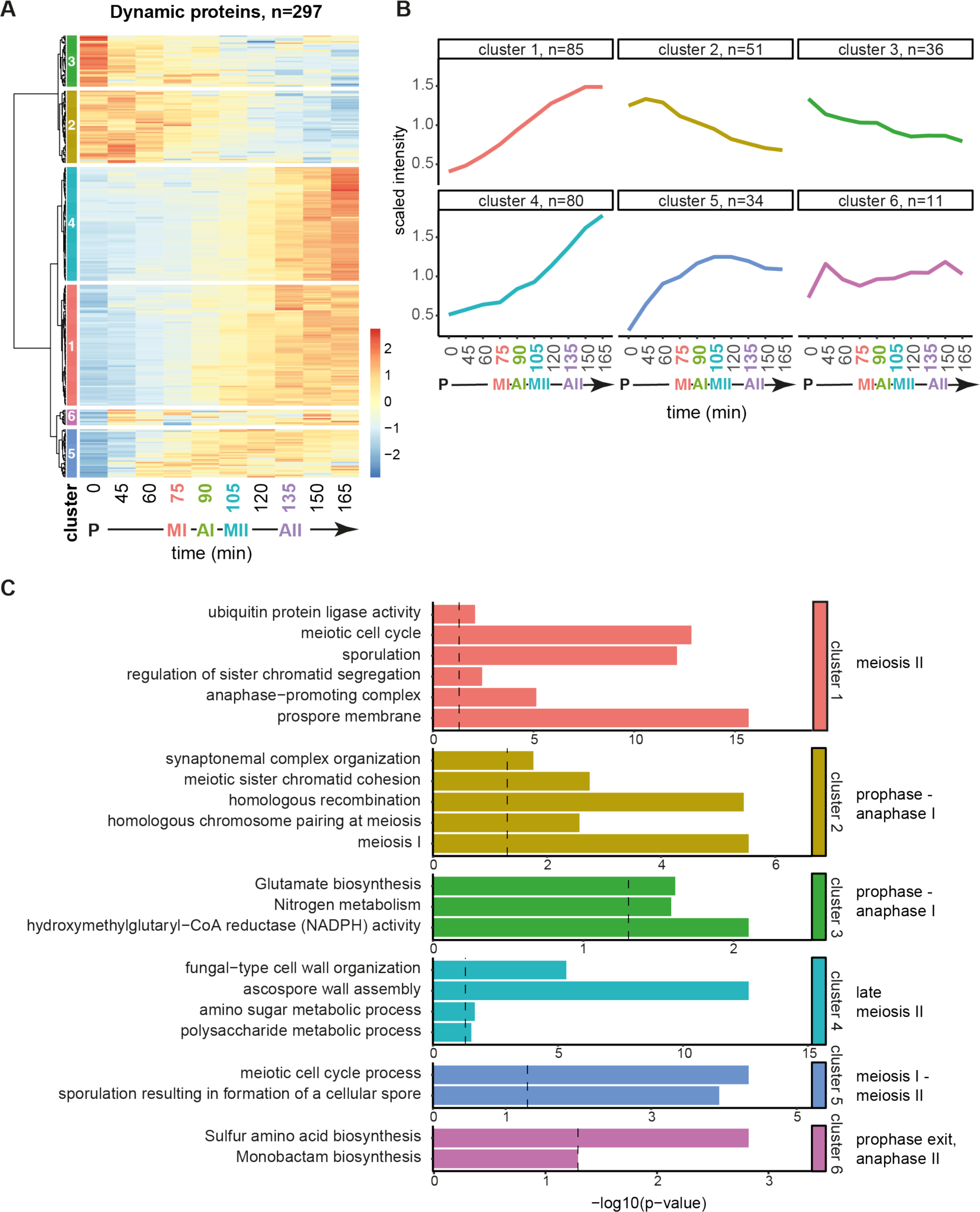
Protein dynamics across a synchronous meiotic division cycle. A. Hierarchical clustering of abundances of dynamic proteins across the timecourse. See Appendix Table S3 for list of proteins included in each cluster. B. Median trend of proteins in each cluster from A. C. GO term enrichment of proteins in each cluster from A and B.

Major cell cycle transitions occur upon anaphase I and II entry where the APC promotes degradation of key cell cycle regulators, including Pds1^securin^ (Cohen-Fix *et al*, 1996; Salah & Nasmyth, 2000). Comparing to their respective metaphase, we observed a significant change in abundance for 54 proteins in anaphase I and 44 proteins in anaphase II, including a decrease in Pds1^securin^ in both cases (Fig EV2A and B). The cell cycle protein, Sgo1, and two APC regulators, Mnd2 and Acm1, (Penkner *et al*, 2005; Oelschlaegel *et al*, 2005; Enquist-Newman *et al*, 2008; Eshleman & Morgan, 2014; Clift *et al*, 2009)) decreased in anaphase I, while the translational repressor Rim4, whose degradation in anaphase II promotes meiotic exit, decreased in anaphase II (Fig EV2B; (Berchowitz *et al*, 2013, 2015; Wang *et al*, 2020)). Consistently, GO terms related to meiosis and the cell cycle were enriched among proteins that decreased in anaphase I and II (Fig EV2C and D), while sporulation and cell wall proteins were among those that increased (Fig EV2C and D). Therefore, despite representing only a small fraction of the proteome, dynamic proteins are involved in key meiotic processes whose function can be predicted by hierarchical clustering. However, the abundance of only a small fraction of proteins changes, indicating that other mechanisms play a major role in orchestrating the meiotic divisions.

### Distinct waves of phosphorylation signatures across the meiotic divisions

To discover the phosphorylations that occur throughout the meiotic divisions we applied the same thresholds used for the protein data, comparing prophase to each of the other timepoints. We identified 1,233 significantly dynamic phosphorylation sites of which the majority (78%) showed increased abundance after prophase, while ∼21% decreased and the remaining 1% of sites were variable depending on the timepoint analysed (Fig EV1B). Hierarchical clustering produced 11 clusters with specific temporal patterns of phosphorylation (Fig 3A) related by common biological processes (Fig 3B). To determine whether these temporal patterns of phosphorylation are driven by particular kinases, we analysed the amino acid sequence surrounding the phospho-sites in each cluster (Fig EV3A). Over 50% of the phospho-sites in cluster 2, where phosphorylation peaks in metaphase I and II, included a proline at the +1 position from the phospho-acceptor residue (Fig 3C). This is the minimal consensus motif ([ST]*P) recognized by Cdc28^Cdk^, suggesting bi-phasic activation of this kinase, consistent with previous studies (Carlile & Amon, 2008). Over 50% of the phosphorylated residues in cluster 3 matched the minimal consensus [DEN]x[ST]*, indicative of a peak of Cdc5^Polo^-dependent phosphorylation in meiosis I (Fig 3C).

**Figure 3.**
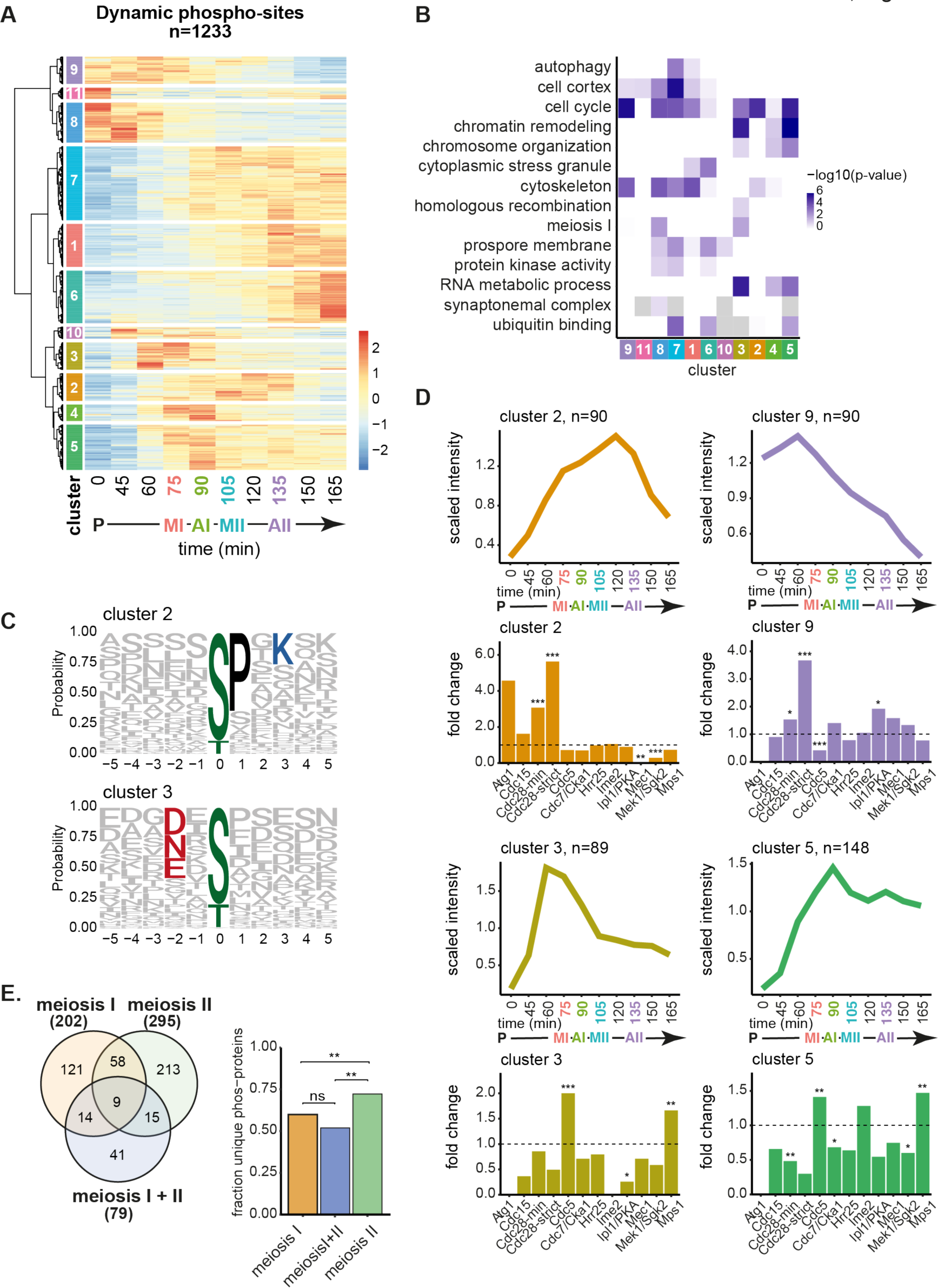
The landscape of phosphorylation across the meiotic divisions. A. Hierarchical clustering of dynamic phospho-sites. See Appendix Table S5 for list of phospho-sites included in each cluster. B. GO term enrichment of clusters. C. Motif logos of selected clusters. D. Median lineplots of selected clusters with kinase motif enrichment analysis bar graph. Asterisks represent p-value from Fisher’s exact test (p<0.001 = ***, p<0.01 = **, p<0.05 = *). See Table EV1 for motif list. E. Venn diagram of meiosis I and meiosis II phospho-proteins grouped by clustering. Bar graph of fraction unique phospho-proteins in each group. Asterisks indicate p-value from Fisher’s exact test (p<0.001 = ***, p<0.01 = **, p<0.05 = *).

To gain broader insight into patterns of kinase activity, we asked whether consensus phosphorylation motifs derived from an *in vitro* analysis using peptide arrays (Mok *et al*, 2010) were enriched in each of the time-resolved clusters. We selected 12 kinases expected to be involved in meiosis and identified clusters that were enriched in consensus motifs for Cdc28^Cdk^, Cdc5^Polo^, Mps1, Ipl1^AuroraB^ and Mek1 (Fig EV3B and C, EV4; Table EV1). The motifs for the major mitotic/meiotic kinases Cdc28^Cdk^ and Cdc5^Polo^ were enriched in several clusters representing distinct waves of phosphorylation (Fig 3D and EV4; Clusters 2-5 and 9). Cdc28^Cdk^ phosphorylation was enriched in prometaphase I (cluster 9), metaphase I (cluster 2) and metaphase II (cluster 2) but significantly depleted from clusters with high phosphorylation in prophase (cluster 8), anaphase I (cluster 4 and 5) or anaphase II and late meiosis (clusters 5 and 6). Cdc5^Polo^/Mps1 kinase-dependent phosphorylation appeared high throughout metaphase and anaphase of both meiotic divisions (clusters 3, 4 and 5). Prophase and prometaphase I clusters (Fig EV4, clusters 8 and 9) were enriched for the Aurora B (Ipl1) consensus, consistent with its role in kinetochore disassembly and biorientation of homologous chromosomes, respectively (Miller *et al*, 2012; Chen *et al*, 2020; Meyer *et al*, 2013, 2015). Similarly, the meiosis-specific Mek1 kinase, which regulates inter-homolog recombination, appeared most active in prophase (Fig EV4, clusters 8 and 11). The meiosis-specific cell cycle regulatory Ime2 kinase, consensus had few matches overall (Fig EV3B and C) but was enriched at prophase exit (cluster 10) and in anaphase I (clusters 4, 5 and 10), though not significantly (Fig EV4).

Focusing on the metaphase I-anaphase I transition, the majority of the 54 phospho-sites which decreased matched either the Cdc5^Polo^ or Ipl1^AuroraB^/PKA consensus, while around half of the 138 sites that increased matched either Cdc5^Polo^, Ipl1^AuroraB^/PKA or the minimal Cdc28^Cdk^ motif (Fig EV5A). At the metaphase II-anaphase II transition, the largest proportion of the 129 decreasing phospho-sites matched the minimal [ST]*P Cdk consensus, while no kinase motif was predominant among the 161 phospho-sites which increased (Fig EV5B). This suggests that the major wave of Cdc28^Cdk^ inactivation occurs at meiosis II and that dynamic Cdc5^Polo^ dependent phosphorylation occurs in meiosis I. However, we found no strong evidence that sites phosphorylated by particular kinases are subject to different rates of dephosphorylation at either anaphase I or II (Fig EV5C and D). The only exception was the strict Cdc28^Cdk1^ consensus ([ST]*Px[KR]), which declined slightly faster in both cases (Fig EV5C and D) and was included in the analysis because the minimal [ST]P motif can also be phosphorylated by other kinases (Mok *et al*, 2010). Furthermore, in contrast to mammalian mitotic exit and during the meiosis I to II transition in starfish (Swartz *et al*, 2021; Holder *et al*, 2020), we also observed no strong preference for dephosphorylation of threonine over serine at either anaphase I or anaphase II (Appendix Fig S2), although it remains possible that there are subtle changes in timing that are below the resolution of our dataset.

It is notable that two clusters of sites that were highly phosphorylated starting in meiosis II (clusters 1, 7) were not significantly enriched for any of the tested motifs (Fig EV4). Similarly, more than half of the phospho-sites increasing after metaphase II did not match the consensus for any of the tested kinases (Fig EV5B). Analysis of the amino acid sequence surrounding the phospho-site in cluster 7 suggested a mixed motif (Fig EV3A), likely due to the activity of multiple kinases on the different substrates. However, cluster 1 phospho-sites were enriched for serine and acidic residues in the surrounding sequence (Fig EV4C). Analysis of the 95 “other” phospho-sites that increased after metaphase II (Fig EV5B) revealed a general enrichment for phospho-acceptor residues and acidic residues in the surrounding sequence (Fig EV5E) which are favoured by the casein kinases (Venerando *et al*, 2014; Mok *et al*, 2010). An attractive candidate kinase for phosphorylating these sites is the Hrr25^CK1^ which is required for spore formation (Argüello-Miranda *et al*, 2017) and increases in abundance during meiosis II (Fig EV5F), though other casein kinases may also play important roles at this stage in meiosis.

Finally, we asked whether the same or different proteins are phosphorylated in meiosis I and II by investigating the overlap between clusters specific to meiosis I (clusters 3,4,9,10), specific to meiosis II (clusters 5,7) or with high phosphorylation during both divisions (cluster 2). This showed that these three groups include mostly unique phospho-proteins and that there is an enrichment of phospho-proteins uniquely present in meiosis II (Fig 3E), suggesting that distinct regulatory mechanisms underlie the two meiotic divisions.

Together, our combined clustering and motif enrichment analysis has identified the temporal order of phosphorylation during the meiotic divisions and predicted the likely kinases responsible. Our data reveal signatures of Ipl1^Aurora^ ^B^ phosphorylation mainly in prophase/prometaphase, distinct waves of Cdc28^Cdk^ and Cdc5^Polo^ kinase phosphorylation in meiosis I and meiosis II and suggest upregulation of Hrr25^CK1^ activity in meiosis II. We also identify patterns of phosphorylation and motifs where the kinase responsible is not easily predictable, which could indicate a role for uncharacterised kinases in key meiotic transitions.

### Proteome changes in *spo13Δ* cells are limited to a small number of key regulators

To determine how two distinct divisions are executed, we analysed the *spo13Δ* mutant which undergoes only a single meiotic division. We generated replicate proteome and phosphoproteome datasets of *spo13Δ* cells synchronously released from prophase as for wild type (Appendix Fig S3). The number of proteins and phosphorylation sites identified and the overall distribution of total protein and phospho-site abundance was equivalent between strains and replicates (Appendix Fig S3 and S4). For comparisons of wild type and *spo13Δ*, we restricted our analysis to proteins identified in at least one timepoint of all 4 experiments and to singly phosphorylated sites that were detected in at least two sequential timepoints with high confidence (localization score > 0.75) of all 4 experiments. Of 3,296 proteins identified in both wild type and *spo13Δ*, we found 414 proteins that were significantly dynamic across the timecourse in either or both strains (Fig EV6A). As a more direct measure of differences between wild type and *spo13Δ*, we performed pair-wise comparisons of protein abundances at matched timepoints to identify proteins which varied by more than 1.5 fold reliably between replicates. This identified 50 proteins which differ in abundance between wild type and *spo13Δ* at any one timepoint, corresponding to 1.5% of the shared wild type-*spo13Δ* proteome (Fig EV7A). Differences included both increases and decreases in abundance and were observed at all stages of meiosis (Fig EV7B), consistent with the wide-ranging phenotypic effects of *spo13Δ.* Proteins that showed altered abundance in *spo13Δ* included key meiotic regulators Pds1^securin^ and Sgo1 which showed reduced and delayed accumulation in *spo13Δ* cells (Fig EV7B and C; cluster 7). Some proteins involved in spore formation (Sps1, Sps22, Spr1) accumulated prematurely while the m6A methyltransferase complex component, Vir1, which is required for the initiation of the meiotic programme (Park *et al*, 2023), was elevated in *spo13Δ* cells (Fig EV7B and C). Regulators Spo12 and Bud14 of the Cdc14 phosphatase, which is required for two meiotic divisions (Kocakaplan *et al*, 2021; Marston *et al*, 2003; Fox *et al*, 2017), were also deregulated in *spo13Δ* cells. Therefore, although overall protein dynamics in *spo13Δ* are similar to wild type, the strict ordering of meiosis I and II events is lost for some key cell cycle and differentiation proteins. This implies that a combination of delayed meiosis I events and premature meiosis II events could explain the mixed single division of *spo13Δ* cells.

### Spo13^Meikin^ influences the dynamics of phosphorylation by major meiotic kinases

Spo13^Meikin^ controls the meiotic programme at least in part through its association with Cdc5^Polo^, but the effect on global phosphorylation remains unknown. We found that 982 phosphorylation sites identified in both wild type and *spo13Δ* were dynamic, meaning that they varied significantly from time zero in one or both strains (Fig EV6B).

As a more direct approach to find the phospho-sites which differ between *spo13Δ* and wild-type, we compared their abundance at each timepoint. This identified 305 phosphorylation sites that were significantly different in abundance, representing ∼13% of the shared phosphoproteome (Fig 4A). Of these sites, the majority (203 or 67%) had reduced phosphorylation in *spo13Δ* versus wild type. To determine whether this could be caused by failure of a particular kinase to phosphorylate these substrates in *spo13Δ*, we asked whether any motifs were over-represented among this group of sites. We used IceLogo (Colaert *et al*, 2009) to compare these 203 sites showing reduced phosphorylation in *spo13Δ* to those showing increased phosphorylation or no change, which showed that asparagine (N) and/or glutamic acid (E) in the -2 position was over-represented (Fig 4B). This matches the canonical Polo kinase recognition motif, [DEN]x[ST]*, and the depletion of phosphorylation on these sites in *spo13Δ* suggests that an important function of Spo13^Meikin^ is to promote Cdc5^Polo^-dependent phosphorylation on at least a subset of its meiotic targets. There was also a group of sites with increased phosphorylation in *spo13Δ* when compared to those that did not change (86 or 28%; Fig 4A), which showed enrichment for arginine (R) in the -3 position, matching the consensus for Ipl1^Aurora^ ^B^ among other basophilic kinases (Fig 4C). This likely indicates repeated rounds of Ipl1^AuroraB^-mediated error correction to reorient incorrect kinetochore-microtubule interactions in *spo13Δ* cells due to the absence of monopolin (Nerusheva *et al*, 2014; Katis *et al*, 2004; Lee *et al*, 2004; Monje-Casas *et al*, 2007).

**Figure 4.**
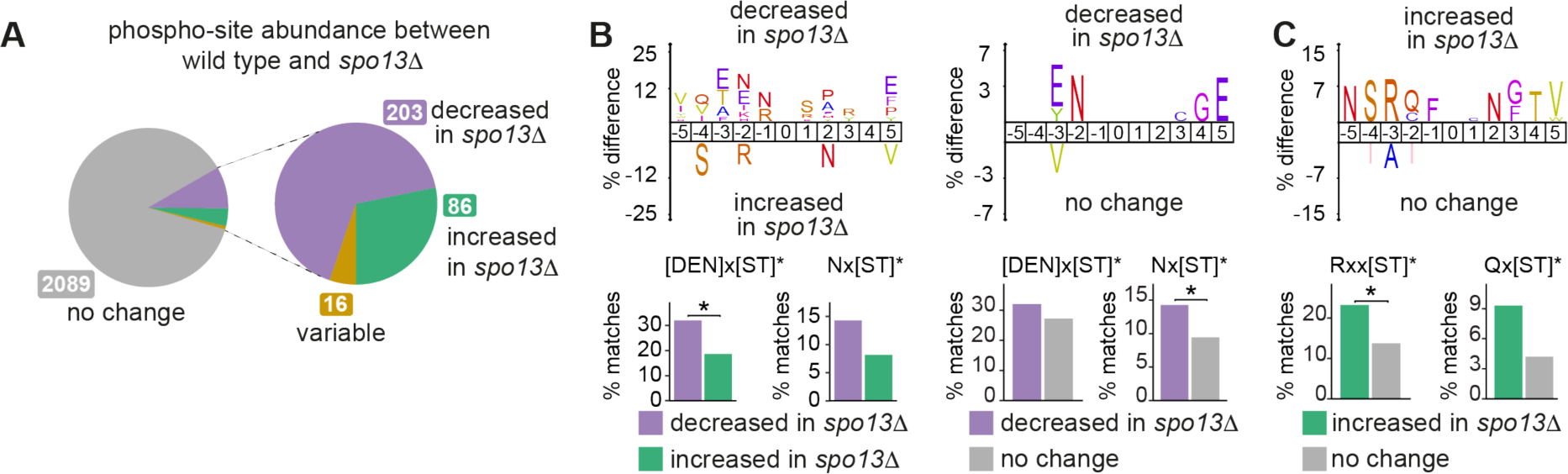
Cdc5^Polo^ kinase phosphorylation is decreased during the meiotic divisions in *spo13Δ* cells. A. Proportion of phospho-sites which significantly vary between wild type and *spo13Δ* at matched time points. See Appendix Table S8 for list of phospho-sites. B. IceLogos showing enrichment of specific residues surrounding the phospho-sites decreased in *spo13Δ.* Bar charts below show percent motif matches in the indicated groups of sites. Asterisks denote p-value from Fishers exact test (p<0.001 = ***, p<0.01 = **, p<0.05 = *). C. IceLogos showing enrichment of specific residues surrounding the phospho-sites increased in *spo13Δ.* Bar charts below show percent motif matches in the indicated groups of sites. Asterisks denote p-value from Fishers exact test (p<0.001 = ***, p<0.01 = **, p<0.05 = *).

To determine whether the differences in phosphorylation between wild type and *spo13Δ* show any temporal trends, we performed hierarchical clustering, which revealed that several groups of sites which peak over the meiotic divisions in wild type are depleted in *spo13Δ* (Fig 5A and B). Cluster 10 showed a peak of phosphorylation in meiosis I in wild type and a delay in phosphorylation in *spo13Δ* and was enriched for the strict Cdc28^Cdk^ motif [ST]*x[KR] (Fig 5C). Interestingly, this cluster included phosphorylation of S36 on the spindle pole body component Spc110 (Fig 5C), which promotes timely mitotic exit (Abbasi *et al*, 2022), and could therefore contribute to the single division meiosis phenotype of *spo13Δ* cells. Cluster 8, which peaks in metaphase I-anaphase I in wild type, but not *spo13Δ*, is strongly enriched for the Cdc5^Polo^ consensus motif (Fig 5D). Among these is a site on the synaptonemal complex component Ecm11 (S169; Fig 5D), which could potentially be a relevant target for Cdc5^Polo^-mediated synaptonemal complex disassembly (Argunhan *et al*, 2017). Cluster 3 is characterised by a sharp rise in phosphorylation after meiosis II which is absent in *spo13Δ* and is enriched for acidic residues typically targeted by Hrr25^CK1^ (Fig 5E). This further strengthens the idea that Hrr25^CK1^-dependent phosphorylation is prevalent in late meiosis II (Fig EV5E and F) and is consistent with a lack of coordinated Hrr25^CK1^ activity in *spo13Δ* cells (Galander *et al*, 2019). Cluster 3 is exemplified by Cdc3-S77, a site within a component of the septin cytoskeleton that is required for efficient spore formation (Heasley & McMurray, 2016), a key function of Hrr25^CK1^ (Argüello-Miranda *et al*, 2017).

**Figure 5.**
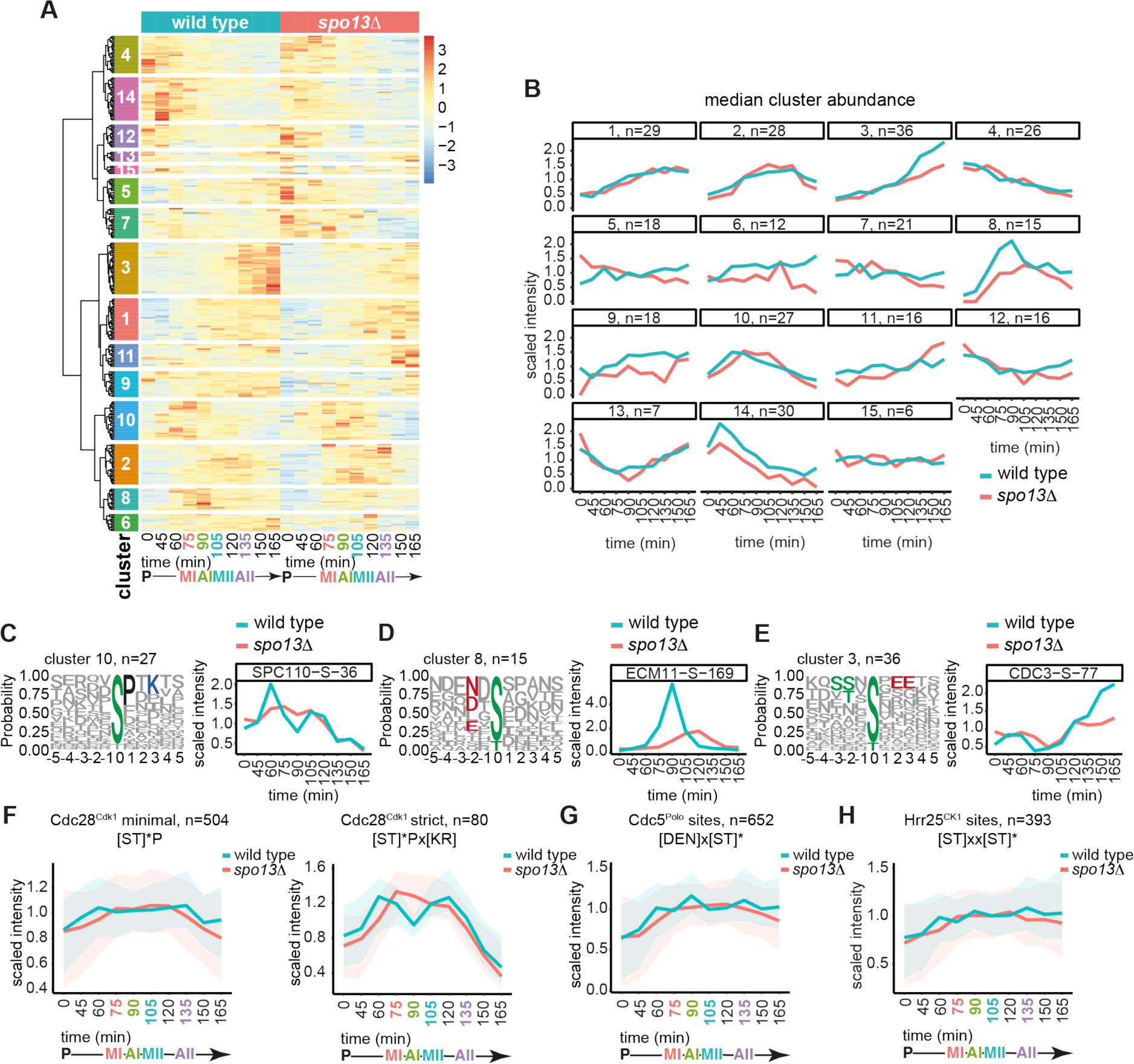
Cdc28^Cdk1^, Cdc5^Polo^, and Hrr25^CK1^ phosphorylation is disrupted in *spo13Δ* cells. A. Hierarchical clustering of the 305 phospho-sites which significantly vary between wild type and *spo13Δ* at matched time points. See Appendix Table S9 for phospho-site identities in each cluster. B. Median line plots of cluster abundances from A. C. Motif logo of cluster 10 contains Cdc28^Cdk1^ consensus, [ST]*Px[KR], matches (left). Abundance of [ST]*x[KR] site Spc110-S36 across the timecourse (right). D. Motif logo of cluster 8 contains Cdc5^Polo^ consensus, [DEN]x[ST]*, matches (left). Abundance of [DEN]x[ST]* site Ecm11-S169 across the timecourse (right). E. Motif logo of cluster 3 contains Hrr25^CK1^ consensus, [ST]xx[ST]* matches as well as acidic or phospho-acceptor residues at -2, +2 and +3 that are recognized by casein kinases (left). Abundance of SSx[ST]* site Cdc3-S77 across the timecourse (right). F. Abundance of phospho-sites matching the Cdc28^Cdk1^ minimal [ST]*P (left) or strict [ST]*x[KR] (right) consensus among sites detected in both replicates of wild type and *spo13Δ* across the timecourse. G. Abundance of phospho-sites matching the Cdc5^Polo^, [DEN]x[ST]*, consensus among sites detected in both replicates of wild type and *spo13Δ* across the timecourse. H. Abundance of phospho-sites matching the Hrr25^CK1^, [ST]xx[ST]*, consensus among sites detected in both replicates of wild type and *spo13Δ* across the timecourse.

To ask whether these effects of *spo13Δ* on the dynamics of phosphorylation by Cdc28^Cdk1^, Cdc5^Polo^ and Hrr25^CK1^ were observable in the broader dataset we plotted the mean scaled abundance of sites matching the consensus for each kinase across the time-course. Only sites detected in both replicates of both wild-type and *spo13Δ* timecourses were included in this analysis (Fig 5F-H). The minimal Cdc28^Cdk1^ consensus increased in abundance from prometaphase I in wild type, declining only after anaphase II, while the strict Cdc28^Cdk1^ consensus revealed a clear bimodal pattern (Fig 5F). In *spo13Δ* both the minimal and strict Cdc28^Cdk1^ consensus accumulated with a delay and decreased prematurely (Fig 5F). The consensus motifs for both Cdc5^Polo^ and Hrr25^CK1^ peaked in prometaphase I (60 min), anaphase I (90 min) and anaphase II (105min) in wild type, but clear peaks were absent in *spo13Δ* (Fig 5G and H). These findings confirm that meiosis I and II are characterised by distinct patterns of phosphorylation and that Spo13 ^Meikin^ is critical to promote this ordering.

In summary, our data suggest that Spo13^Meikin^ promotes phosphorylation of a group of substrates by Cdc5^Polo^ kinase specifically in meiosis I and likely affects Hrr25^CK1^ function in late meiosis II. Additionally, we find that the presence of Spo13^Meikin^ limits Ipl1^Aurora^ ^B^ kinase-dependent phosphorylation in early meiosis and promotes two consecutive peaks of Cdc28^Cdk1^ activity.

### Spo13^Meikin^ promotes Cdc28^Cdk1^ and Cdc5^Polo^ consensus phosphorylation at metaphase I

Spo13^Meikin^ is expected to exert its key function in metaphase I prior to its degradation in anaphase I (Sullivan & Morgan, 2007). To gain deeper insight into the effect of Spo13^Meikin^ and to understand the importance of Cdc5^Polo^ binding, we generated metaphase I proteomic and phosphoproteomic datasets of quadruple wild-type and triplicate *spo13Δ* and *spo13-m2* cultures (*spo13-m2* carries a mutation in the motif that binds the Polo Binding Domain (PBD) of Cdc5^Polo^ (Matos *et al*, 2008)). Metaphase I arrest and the even distribution of protein and phosphorylation sites were confirmed and 3,960 proteins and 6,927 phosphorylation sites were identified in at least 3 replicates of wild type, *spo13Δ* and *spo13-m2* strains (Appendix Fig S5, bottom row).

Between wild type and *spo13Δ* or wild type and *spo13-m2*, 115 proteins (1.9%) or 34 proteins (0.9%), respectively, showed significantly different abundance at metaphase I (Fig 6A and B; Appendix Fig S6). Consistent with the timecourse (Fig EV7), the mitotic exit network factor Spo12 was less abundant in *spo13Δ* at metaphase I (Appendix S6). The m6A methyltransferase complex (MIS) complex (Vir1, Kar4 and Mum2) and the nuclear envelope lipid synthesis factor Nem1, which were less abundant in *spo13Δ* at metaphase I (Appendix S6), were also significantly changed between strains in the timecourse, although the overall direction of change was opposite in that context. This suggests that Spo13^Meikin^ has effects on protein abundance prior to anaphase I and raises the interesting possibility that Spo13 controls gene expression. We note that *spo13-m2* has a more modest effect on protein abundance than *spo13Δ* (Fig 6A and B), suggesting that Spo13^Meikin^ may affect protein abundance independently of Cdc5^Polo^ binding. Indeed, Cdc5^Polo^ accumulates only after prophase I exit (Sourirajan & Lichten, 2008), so any prior Spo13^Meikin^ functions would be expected to be mediated independently. However, an important caveat is that Spo13-m2 retains residual Cdc5^Polo^ binding (Matos *et al*, 2008) which could account for its more modest effects.

**Figure 6.**
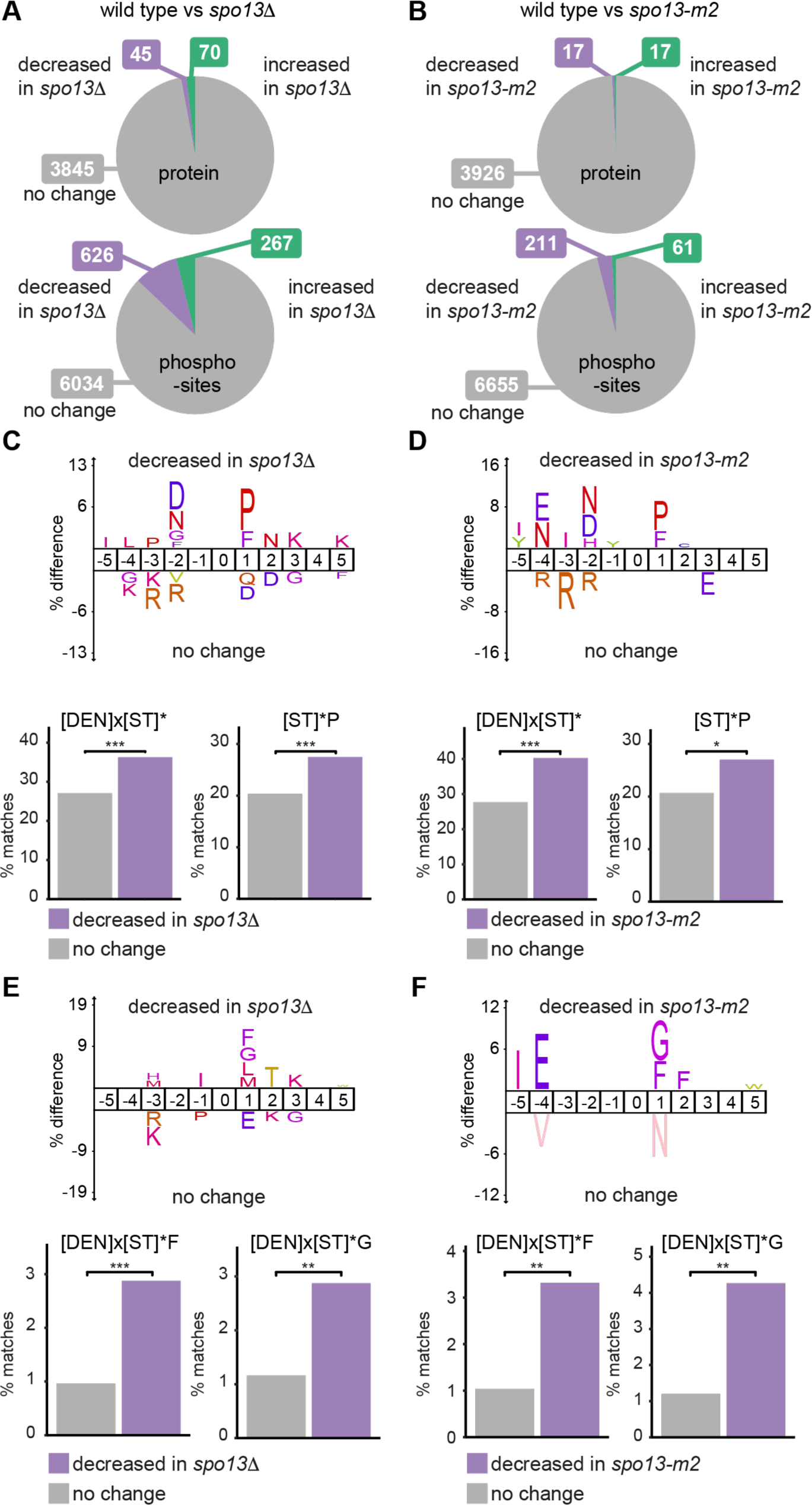
At metaphase I, Cdc5^Polo^ kinase phosphorylation is decreased in *spo13Δ* and *spo13-m2* cells. A. Proportion of proteins and phospho-sites which significantly vary between wild type and *spo13Δ* in metaphase I arrested cells. See also Appendix Tables S10 and S12. B. Proportion of proteins and phospho-sites which significantly vary between wild type and *spo13-m2* in metaphase I arrested cells. See also Appendix Tables S11 and S13. C. IceLogo of motifs enriched surrounding phospho-sites decreased in *spo13Δ* in metaphase I arrested cells. Bar charts below show percent motif matches in the indicated groups of sites. Asterisks denote p-value from Fishers exact test (p<0.001 = ***, p<0.01 = **, p<0.05 = *). D. IceLogo of motifs enriched surrounding phospho-sites decreased in *spo13-m2* in metaphase I arrested cells. Bar charts below show percent motif matches in the indicated groups of sites. Asterisks denote p-value from Fishers exact test (p<0.001 = ***, p<0.01 = **, p<0.05 = *). E. IceLogo of motifs enriched surrounding [DEN]x[ST]* phospho-sites decreased in *spo13Δ* in metaphase I arrested cells. Bar charts below show percent motif matches in the indicated groups of sites. Asterisks denote p-value from Fishers exact test (p<0.001 = ***, p<0.01 = **, p<0.05 = *). F. IceLogo of motifs enriched surrounding [DEN]x[ST]* phospho-sites decreased in *spo13-m2* in metaphase I arrested cells. Bar charts below show percent motif matches in the indicated groups of sites. Asterisks denote p-value from Fishers exact test (p<0.001 = ***, p<0.01 = **, p<0.05 = *).

Compared to wild type at metaphase I, 893 and 272 phospho-sites were significantly different in *spo13Δ* and *spo13-m2*, respectively (Fig 6A and B). As in the timecourse experiments (Fig 4A), the dominant trend was for reduced phosphorylation in *spo13Δ* compared to wild type (626/893 or 70% of significantly different sites). Although fewer significantly different phosphorylation sites were identified in the *spo13-m2* mutant compared to wild type, the fraction that decreased was very similar to *spo13Δ* (211/272 or 78%). Motif analysis among sites showing reduced phosphorylation in *spo13Δ* or *spo13-m2* versus wild type revealed an enrichment for aspartic acid /asparagine at -2 and proline at +1, consistent with reduced Cdc5^Polo^ and Cdc28^Cdk1^ phosphorylation in the absence of functional Spo13^Meikin^ (Fig 6C and D).

### Identification of a Spo13^Meikin^ - Cdc5^Polo^ consensus motif

In contrast to *spo13Δ*, depletion of Cdc5^Polo^ leads to metaphase I arrest (Lee & Amon, 2003; Clyne *et al*, 2003), indicating that Spo13^Meikin^ must control phosphorylation of only a subset of Cdc5^Polo^ target sites. Indeed, only ∼10-12% of Cdc5^Polo^-motif matching sites had decreased phosphorylation in *spo13Δ* in both the metaphase I arrest and the timecourse experiments (Fig EV8A and B). This raises the question of how Spo13^Meikin^ enhances Cdc5^Polo^ kinase phosphorylation for only a subset of substrates. We hypothesised that Spo13^Meikin^ may direct Cdc5^Polo^ to a subset of targets containing a specific motif. To test this idea, we searched for sub-motifs among the differentially phosphorylated Cdc5^Polo^ motif-matching sites,[DEN]x[ST]*, in wild type versus *spo13Δ* or *spo13-m2* in metaphase I and discovered a preference for phenylalanine (F) or glycine (G) in the +1 position among sites with reduced phosphorylation in either *spo13Δ* or *spo13-m2* compared to wild type (Fig 6E and F). This suggests that Spo13^Meikin^ may specifically or preferentially promote Cdc5^Polo^-directed phosphorylation of substrates carrying the novel motif [DEN]x[ST]*[FG].

### Spo13^Meikin^ -Cdc5^Polo^ -dependent phosphorylation occurs on chromosome related proteins

To understand whether phospho-proteins regulated by Spo13^Meikin^ at metaphase I have shared functions, we performed GO term enrichment analysis. Among proteins with decreased phosphorylation in *spo13Δ* or *spo13-m2* vs wild type in metaphase I arrest, the terms “chromatin remodelling” and “RNA biosynthetic process” were enriched (Fig EV8C). For proteins with increased phosphorylation in *spo13Δ*, cytoskeleton-related terms “prospore membrane” and “cytoskeleton organization” were most enriched (Fig EV8C). To look more specifically at proteins regulated by Cdc5^Polo^ phosphorylation, we restricted the GO term analysis to phospho-sites matching the canonical Cdc5^Polo^ motif, [DEN]x[ST]*, among sites with differential phosphorylation in *spo13*Δ in metaphase I arrest. Reflecting the primary effect of Spo13^Meikin^ on the phosphorylation of Cdc5^Polo^ substrates, nearly the same terms were enriched (Fig 7A). Further restricting our analysis to the 74 proteins phosphorylated at metaphase I on the Spo13^Meikin^-Cdc5^Polo^ consensus [DEN]x[ST]*F identified above, GO terms enriched included “meiotic chromosome separation”, “chromosome organization” and “chromatin binding” (Fig 7B). The majority (57%) of these proteins phosphorylated on [DEN]x[ST]*F in the metaphase I arrest were also phosphorylated in our timecourse analysis (42/74) (Fig 7C), consistent with Spo13^Meikin^ exerting its effects in metaphase I and providing confidence that these sites represent functionally important substrates.

**Figure 7.**
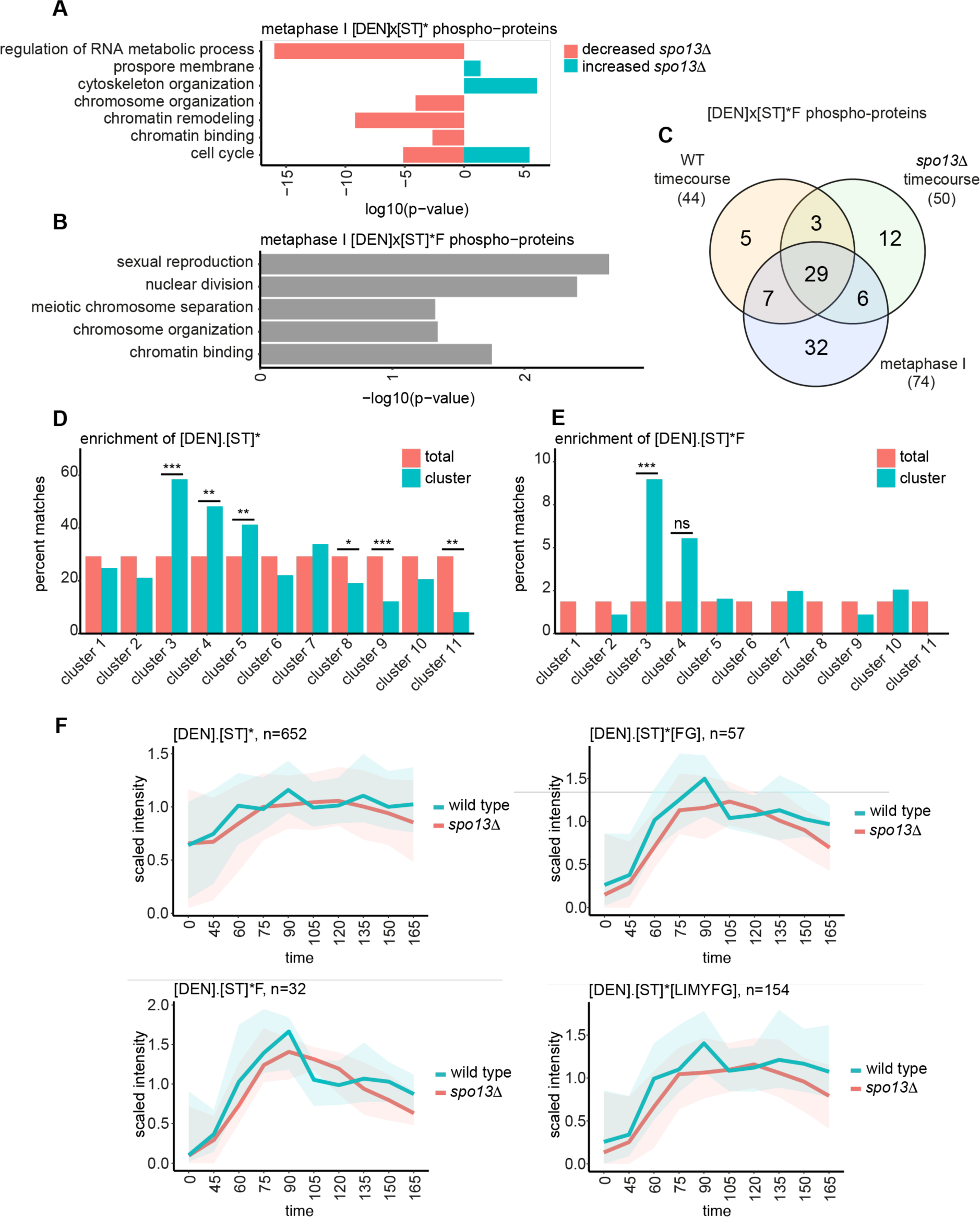
Spo13^Meikin^ promotes [DEN]x[ST]*F phosphorylation in metaphase I. A. Bar graph of selected GO terms enriched among proteins with phosphorylated [DEN]x[ST]* sites with significantly different abundance between wild type and *spo13Δ* in the metaphase I arrest dataset. B. Bar graph of selected GO terms enriched among proteins with phosphorylated [DEN]x[ST]*F sites in the metaphase I arrest dataset. C. Proportion of phospho-proteins with phosphorylated [DEN]x[ST]*F motifs detected in the wild type timecourse, *spo13Δ* timecourse, and both wild type and *spo13Δ* in the metaphase I arrest experiment. D. Percent [DEN]x[ST]* motif matches in the indicated groups of sites. Clusters refer to wild type phospho-site clustering in Fig 3 and EV4. Asterisks denote p-value from Fisher’s exact test (p<0.001 = ***, p<0.01 = **, p<0.05 = *). E. Percent [DEN]x[ST]*F motif matches in the indicated groups of sites. Clusters refer to wild type phospho-site clustering in Fig 3 and EV4. Asterisks denote p-value from Fisher’s exact test (p<0.001 = ***, p<0.01 = **, p<0.05 = *). F. Abundance of phospho-sites matching the indicated motifs among sites detected in both replicates of wild type and *spo13Δ* across the timecourse.

### Spo13^Meikin^ directs Cdc5^Polo^ to novel consensus motif [DEN]x[ST]*F in meiosis I

Spo13^Meikin^ and Cdc5^Polo^ co-exist at prometaphase/metaphase I, after which Spo13^Meikin^ is degraded. Therefore, if Spo13^Meikin^-Cdc5^Polo^ is responsible for phosphorylation of [DEN]x[ST]*F, it is expected to be maximal at metaphase I. To test this idea, we returned to the timecourse experiment, where we had noticed a slight preference for +1F among sites highly phosphorylated specifically at metaphase I in wild type (Fig 3C, cluster 3). We directly tested for the enrichment of the motif [DEN]x[ST]*F among each of the 11 phospho-site clusters from the wild type timecourse and found that among the three clusters enriched for the canonical Cdc5^Polo^ motif [DEN]x[ST]* (Fig 7D), only cluster 3 was significantly enriched for the novel [DEN]x[ST]*F motif (Fig 7E). Interestingly, cluster 3 is the only Cdc5^Polo^ motif-enriched cluster that is specific to metaphase I (Fig 3D and EV4). This strongly suggests that phosphorylation of the novel Spo13^Meikin^-dependent [DEN]x[ST]*F motif occurs predominantly in metaphase I.

To investigate the timing of phosphorylation of these sites in more detail in wild type and *spo13Δ*, we plotted the average abundance of all sites matching the [DEN]x[ST]*F, [DEN]x[ST]*G and [DEN]x[ST]*[FG] motifs across the timecourse (Fig 7F and EV8D). Since previous studies have mentioned a slight preference for Polo kinase to phosphorylate substrates with hydrophobic or aromatic residues at +1 (Mok *et al*, 2010; Santamaria *et al*, 2011), we also extended our analysis to the motif [DEN]x[ST]*[LIMYFG]. In wild type, all motifs showed two peaks of abundance in metaphase I-anaphase I (75-90min) and in metaphase II (135 min) (Fig 7F, EV8D). With the exception of [DEN]x[ST]*G (Fig EV8D), these peaks were more pronounced for all extended motifs, compared to the canonical [DEN]x[ST]* motif. Interestingly, phosphorylation of [DEN]x[ST]*F and to a lesser extent [DEN]x[ST]*[FG], was increased in meiosis I relative to meiosis II (Fig 7F), indicating that F in the +1 position is the best predictor of meiosis I-specific phosphorylation of the Cdc5^Polo^ consensus.

Consistently, in *spo13Δ*, the greatest deviation from the wild type trend was observed for Cdc5^Polo^ motifs with F in the +1 position. Phosphorylation of the canonical [DEN]x[ST]* motif shows relatively modest changes in timing in *spo13Δ* compared to wild type (Fig 7F). However, for motifs including +1F, the pronounced meiosis I peak observed in wild type was lost in *spo13Δ* (Fig 7F). This suggests that Spo13^Meikin^ ensures that Cdc5^Polo^ sites with hydrophobic residues, particularly phenylalanine, at +1 are preferentially phosphorylated at metaphase I. Overall, these data identify a novel motif [DEN]x[ST]*F that is specifically phosphorylated in meiosis I by a specialized kinase, Spo13^Meikin^-Cdc5^Polo^.

## Discussion

Here we describe the time-resolved global proteome and phosphoproteome of the meiotic divisions, providing unprecedented insight into the protein and phosphorylation changes that govern this specialized cell cycle. Our datasets provide a valuable resource to discover the mechanisms underlying the regulation of key meiotic processes including kinetochore monoorientation, cohesin protection, regulation of the meiosis I to II transition, spindle pole body duplication, organelle partitioning and gametogenesis. While protein changes were limited to a small number of key regulators, we found that distinct groups of phosphorylations characterised each stage of the meiotic divisions. Matching phosphorylations to consensus motifs revealed that Cdc28^Cdk^ and Cdc5^Polo^ each direct two waves of phosphorylation, corresponding to meiosis I and II. Our data also suggest that while phosphorylation by Cdc28^Cdk^ declines in anaphase I, Cdc5^Polo^ is active in anaphase I, and that a distinct set of proteins are phosphorylated in meiosis I and meiosis II. We found that Spo13^Meikin^ is required for a subset of Cdc5^Polo^-directed phosphorylation in meiosis I, Hrr25^CK1^ casein kinase phosphorylation in meiosis II and for two waves of Cdc28^Cdk^ phosphorylation. Analysis of metaphase I-arrested cells identified a specific motif, [DEN]x[ST]*F, which is enriched in the wild type phosphoproteome at this time but strongly depleted from *spo13Δ* and *spo13-m2* cells. Therefore, by redirecting Cdc5^Polo^ to specific substrates containing the [DEN]x[ST]*F motif at metaphase I, Spo13^Meikin^ establishes the meiotic programme.

### Two waves of Cdk phosphorylation in meiosis

A longstanding question is how cells progress through the meiosis I to meiosis II transition without resetting replication origins. The prevailing hypothesis is that Cdks retain activity towards certain substrates at this transition, however, this has been challenging to address because of poor synchrony in the meiotic progression coupled with the short time interval between anaphase I and metaphase II. Nevertheless, accumulating evidence indicates that Cdks are at least partially inactivated between meiosis I and II (Furuno *et al*, 1994; Iwabuchi *et al*, 2000; Swartz *et al*, 2021; Carlile & Amon, 2008). A second hypothesis is that the activity of a distinct kinase could maintain some phosphorylation at this transition. In support of this idea, Ime2 kinase is at least partially active as cells transition into meiosis II (Berchowitz *et al*, 2013) (see also Fig EV4) and the sites it phosphorylates are resistant to dephosphorylation by the Cdc14 phosphatase (Holt *et al*, 2007). However, to date only a handful of *in vivo* Ime2 substrates have been identified (Berchowitz *et al*, 2013) and its consensus motif partially overlaps with that of many basophilic kinases (Holt *et al*, 2007; Mok *et al*, 2010). Consequently, very few *bone fide* Ime2-specific motif, RPx[ST]*, phospho-sites were found in our dataset (Fig EV3B), precluding further conclusions on the extent of Ime2 activity at the global level. However, a third, not mutually exclusive, hypothesis posits that selective dephosphorylation of substrates could temper a loss of kinase activity at the meiosis I to II transition. In support of this idea, we discovered that strict Cdc28^Cdk1^ consensus sites fall substantially in abundance at meiosis I exit, while the minimal Cdc28^Cdk1^ consensus phospho-sites do not (Fig 5F). This suggests that phosphatases active at the meiosis I to II transition could have preference for Cdc28^Cdk1^ sites with lysine or arginine in the +2 position, while sites with a different residue at +2 are protected from dephosphorylation. Directed mechanistic studies are required to test these ideas and also to understand how Cdks are permitted to rise again for entry into meiosis II.

### Spo13^Meikin^ alters Cdc5^Polo^ substrate specificity

We propose that Spo13^Meikin^ directs Cdc5^Polo^ preferentially to substrates with a modified consensus with phenylalanine in the +1 position [DEN]x[ST*]F, which we refer to as the Spo13 ^Meikin^ -Cdc5 ^Polo^ consensus phosphorylation motif. There are three lines of evidence for this proposal. First, this motif was strongly depleted from the *spo13Δ* and *spo13-m2* metaphase I phosphoproteome. Second, in the wild-type time course experiment, among the clusters where the Cdc5^Polo^ consensus was enriched, only those with high abundance in metaphase I showed enrichment for the F at +1. Third, abundance of [DEN]x[ST]*F reached a clear peak of metaphase I in wild type, but not in *spo13Δ*. The fact that Spo13 is degraded in anaphase I and does not reaccumulate in meiosis II (Sullivan & Morgan, 2007; Katis *et al*, 2004), together with our finding that [DEN]x[ST*]F phosphorylation peaks at metaphase I in wild type further indicates that Spo13^Meikin^-Cdc5^Polo^ is responsible for this phosphorylation.

How might Spo13^Meikin^ influence the substrate choice of Cdc5^Polo^? The question is particularly intriguing since Spo13^Meikin^ contains an [S][ST]*[PX] motif which is required for binding to Cdc5^Polo^ (Matos *et al*, 2008). This suggests that Cdc5^Polo^ uses its Polo Binding Domain (PBD) to bind to phosphorylated Spo13^Meikin^ in the same way it normally binds to substrates (Elia *et al*, 2003). Since the usual mode of substrate binding is blocked, Spo13-Cdc5^Polo^ must therefore use another mechanism to target substrates. Potentially, Spo13^Meikin^ itself targets Spo13^Meikin^-Cdc5^Polo^ to substrates. Indeed, Spo13, as well as fission yeast Moa1 and mouse Meikin all recruit Polo kinase to kinetochores, which in the case of Moa1 and Meikin is through a direct Meikin-CENP-C interaction, however CENP-C itself may not be the Polo relevant substrate (Kim *et al*, 2015; Galander *et al*, 2019; Ma *et al*, 2021). There is also precedent for Cdc5^Polo^ using a different surface of its PBD to associate with binding partners, an interaction that can occur coincidentally with binding to a substrate in the canonical manner (Chen & Weinreich, 2010; Almawi *et al*, 2020). However, how Spo13^Meikin^ binding would alter the substrate preference of Cdc5^Polo^ is unclear. Further understanding of how Spo13^Meikin^ directs Cdc5^Polo^ awaits structural and biochemical studies.

### The phosphorylation events that programme homolog segregation

What are the functional consequences of Spo13^Meikin^ re-wiring the phosphoproteome? We find that chromosome and cell cycle-associated proteins are enriched within the Spo13^Meikin^-Cdc5^Polo^ phosphoproteome. This is consistent with both the documented roles of Spo13^Meikin^ in sister kinetochore monoorientation, cohesion protection and the execution of two meiotic divisions (Lee *et al*, 2004; Katis *et al*, 2004) and the fact that at least a fraction of Spo13^Meikin^ is localized to chromosomes (Galander *et al*, 2019). Our study therefore provides a starting point for future investigations to delineate the molecular mechanisms underlying the functional consequences of Spo13^Meikin^-Cdc5^Polo^-dependent phosphorylation. We note, however, that Cdc5^Polo^ also has meiotic functions independent of Spo13^Meikin^, implying that only a fraction of cellular Cdc5^Polo^ is associated with Spo13^Meikin^ and raising the possibility that Spo13^Meikin^ is rate-limiting. The observation that over-expression of Spo13^Meikin^ blocks cells in mitotic metaphase (Shonn *et al*, 2002; Lee *et al*, 2002; McCarroll & Esposito, 1994; Maier *et al*, 2021) could suggest titration of Cdc5^Polo^ away from key targets, in support of this idea. Understanding the balance between free and Spo13^Meikin^-bound Cdc5^Polo^ will be an important avenue of future investigation.

Although meiotic errors are frequent, causing fertility issues and developmental disorders in humans, the molecular basis of meiosis is much less understood than mitosis, in part due to the scarcity of meiotic material. Our proteomics and phospho-proteomics study in budding yeast has provided a rich dataset that can help to bridge this gap to discover key molecular mechanisms that could be relevant for human fertility.

## Methods

### Yeast strains

Yeast strains used in this study were derivatives of SK1 and are listed in Appendix Table S1. *pCLB2-CDC20* (Lee & Amon, 2003), *pGAL1-NDT80*, *pGPD1-GAL4.ER* (Benjamin *et al*, 2003) and *spo13-m2* (Matos *et al*, 2008) were described previously.

### Immunofluorescence

Meiotic spindles were visualised by indirect immunofluorescence as described in (Barton *et al*, 2022). A rat anti-tubulin primary antibody (AbD serotec) at 1:50 dilution and an anti-rat FITC conjugated secondary antibody (Jackson Immunoresearch) at 1:100 dilution were used.

### Meiotic prophase block-release timecourse

Cells were induced to undergo meiosis as described by (Barton *et al*, 2022) and *pGAL-NDT80* prophase block-release experiments were performed as outlined in (Carlile & Amon, 2008). Briefly, strains were patched from -80 stocks to YPG agar (1% Bacto yeast extract, 2% Bacto peptone, 2.5% glycerol, 0.3 mM adenine, 2% agar) plates. After ∼16h, cells were transferred to 4% YPDA agar (1% Bacto yeast extract, 2% Bacto peptone, 4% glucose, 0.3 mM adenine, 2% agar) at 30°C. After 8-16h, cells were inoculated into YPDA media and grown for 24 h at 30°C with shaking at 250 rpm. Next, BYTA (1% Bacto yeast extract, 2% Bacto tryptone, 1% potassium acetate, 50 mM potassium phthalate) cultures were prepared to OD_600_=0.3 and grown at 30°C with shaking at 250rpm overnight (∼16 h). The next morning, cells were washed twice in sterile water and resuspended in sporulation medium at OD_600_=2.0. After 6 h in sporulation media, 1 μM β-estradiol was added to release cells from prophase and samples were collected for immunofluoresence every 15 min until 180 min, and then every 30 min for a further hour. For preparation of protein extracts for mass spectrometry, 10 ml samples were collected at time 0 and at 45-165 minutes after prophase release by centrifugation at 3,000 rpm for 3 min and the supernatants removed. Cells were resuspended in 5% TCA and incubated on ice for ∼10 minutes. Next the samples were centrifuged again for 3 minutes at 3,000 rpm and the supernatant removed. The samples were transferred to 2 ml Fastprep tubes (MP biomedicals) and spun in a microcentrifuge at 13.2k rpm for 1 min. The supernatant was removed by aspiration and the cell pellets drop frozen in liquid nitrogen before being stored at -80°C before further processing.

### Sample preparation for Tandem Mass Tag (TMT) mass spectrometry

Cell pellets stored at -80°C were placed on ice and then 500 μl ice-cold acetone was added and the samples were vortexed for a few seconds and placed at -20°C for ∼1 hour while urea lysis buffer (8M urea, 75 mM NaCl, 50mM HEPES pH 8.0) was prepared. Samples were spun in a chilled 4°C microcentrifuge at 13.2krpm for 10 minutes before the supernatants were removed and the pellets were dried in the hood for 10 min. Each sample was then resuspended in 200 μl freshly prepared 8M urea lysis buffer, supplemented with 2mM beta glycerophosphate, 1mM Na pyrophosphate, 5mM NaF, 2mM AEBSF, 2X CLAAPE (10ug/ul each of Chymostatin, Leupeptin, Antipain, Aprotinin, Pepstatin A, E-64 protease inhibitor), 0.8 mM NaVO4, 0.2uM microcystin-LR, and 1X Roche protease inhibitor cocktail., by pipetting up and down. Silica beads were added and cells were lysed by bead-beating in a FastPrep (MP Biomedicals) machine at 4C, with 4x 45 s rounds of lysis and 2 mins on ice in between each round. Cell lysates were separated from beads by poking a hole in the bottom of each tube with a red-hot needle and placing the tube on top of a new Fastprep tube and briefly spinning them for ∼20 s in a centrifuge. Next, the samples were spun for 15 minutes at 7k rpm in a chilled 4°C microcentrifuge and the clarified lysate supernatants were transferred to protein lo-bind Eppendorf tubes. A BCA assay (Pierce) was carried out, following the manufacturer’s instructions, to determine protein concentration and 400 μg protein from each sample was processed further.

Each 400 μg protein sample was reduced by adding 5 mM DTT and incubating at 37°C for 15 min with gentle shaking at 500 rpm in an Eppendorf shaker/incubator. Next, samples were alkylated in 10 mM iodoacetamide (IAA) for 30 min in the dark. Then, 15 mM DTT was added and samples were incubated for a further 15 min in the dark. Finally, 5x volume of ice-cold acetone was added to each sample, briefly vortexed, and stored at -20°C overnight (∼16 h).

The next day, samples were spun in a chilled 4°C microcentrifuge at 8,000 g for 10 min, the supernatant removed and the pellets dried in the hood for 10 min. Samples were resuspended in 90 μl of 100 mM TEAB (Thermo Scientific). Samples were digested with 10 μg Trypsin in two rounds to facilitate digestion, with 5 μg Trypsin added and samples incubated 37°C for 4 h, then the remaining 5 μg Trypsin added and samples mixed and incubated 37°C overnight (16h). The next morning, samples were clarified by spinning in a microcentrifuge at 4k rpm for 10 min at room temperature and the supernatant removed to fresh tubes. The final volumes were brought to 100 μl in 100 mM TEAB. Each 0.8 mg TMT reagent (Thermo scientific) was resuspended in 41 μl acetonitrile, mixed with the peptide sample, and the tubes were gently shaken at 400 rpm at 25°C for 1 h. Reactions were quenched with 50 mM Tris pH8.0 and shaken at 400 rpm, 25°C for 15 min. The 10 samples were then combined together in one tube and acidified to pH <3 with formic acid, dried in a vacuum centrifuge and stored at -80°C. The sample was resuspended in 0.1% formic acid, pH <3 and then desalted using a 500 mg C18 SepPak cartridge (Waters). To remove free TMT from samples, an extra wash with 0.5% acetic acid was added to the protocol and elution was done in 70% acetonitrile, diluted in 0.2% formic acid.

Sample was resuspended in 10 mM ammonium formate pH 9.0 and fractionated by high pH reversed-phase (HPRP) chromatography using an Ultimate 3000 HPLC (Thermo Fisher Scientific). Peptides were loaded onto a XBridge Peptide BEH C18 130Å 3.5μm 4.6×150mm column (Waters) and eluted at 1ml/min using a constant 10mM ammonium formate pH 9.0 and a multistep gradient from 2% to 50% acetonitrile over 10 min, 50% to 80% gradient for a further 1 min, followed by an 80% column wash and re-equilibration. 72 fractions were collected at 8.5 s intervals and concatenated into 12 fractions and dried.

∼5% of each fraction was reserved as the non-phospho-enriched (“N”). The remaining 95% was subjected to phospho-peptide enrichment (“PE”) using magnetic Ti4+ IMAC beads (MagReSyn).

Each fraction was phospho-enriched by mixing peptides with 1 mg of MagReSyn Ti-IMAC beads in load buffer, which consisted of 80% acetonitrile, 5% trifluoroacetic acid (TFA) and 5% glycolic acid, for 20 min at 25°C. Beads were washed for 2 min with 80% acetonitrile +1% TFA. This was followed by two 2 min washes in 10% acetonitrile + 0.2% TFA. Phosphorylated peptides were eluted for 15 min twice in 1% ammonium hydroxide. This was followed by a second elution for 1 h in 1% ammonium hydroxide 50% acetonitrile.

### Mass spectrometry

Phospho-peptide enriched samples and reserved non-phospho-enriched samples were then desalted using C18 stage tips (Rappsilber *et al*, 2003) and eluted in 40 μL of 80% acetonitrile in 0.1% TFA and concentrated down to 1 μL by vacuum centrifugation (Concentrator 5301, Eppendorf, UK). They were then prepared for LC-MS3 analysis by diluting it to 5 μL by 0.1% TFA. All fractions from every experiment were injected on Orbitrap Fusion™ Lumos™ Tribrid^TM^ mass spectrometer, coupled on-line, to an Ultimate 3000 HPLC (Dionex, Thermo Fisher Scientific, UK). Peptides were separated on a 50 cm (2 µm particle size) EASY-Spray column (Thermo Scientific, UK), which was assembled on an EASY-Spray source (Thermo Scientific, UK) and operated constantly at 50°C. Mobile phase A consisted of 0.1% formic acid in LC-MS grade water and mobile phase B consisted of 80% acetonitrile and 0.1% formic acid. Peptides were loaded onto the column at a flow rate of 0.3 μL min^-1^ and eluted at a flow rate of 0.25 μL min^-1^ according to the following gradient: 2 to 40% mobile phase B in 120 min and then to 95% in 11 min. Mobile phase B was retained at 95% for 5 min and returned back to 2% a minute after until the end of the run (160 min). Survey scans were recorded at 120,000 resolution (scan range 380-1500 m/z) with an ion target of 4.0e5, and injection time of 50ms. MS2 was performed in the ion trap at a turbo scan mode, with ion target of 2.0E4 and CID fragmentation with normalized collision energy of 35, Q activation parameter at 0.25 and CID activation time at 10ms. The isolation window in the quadrupole was 0.7 Thomson. Only ions with charge between 2 and 7 were selected for MS2. Dynamic exclusion was set at 70 s. MS3 scans were performed in the orbitrap, with the number SPS precursors set to 5. The isolation window was set at 2, and the resolution at 50,000 with scan range 100-150m/z. HCD fragmentation (Olsen *et al*, 2007) was performed with normalized collision energy of 65.

### MaxQuant conditions

All 4 TMT10 time-course experiments, including N and PE samples and 24 fraction samples total per TMT10 experiment, were analysed together in a single MaxQuant (MQ) (Cox & Mann, 2008) session. For the metaphase I arrest experiment, N and PE samples were also analysed in the same MQ session. The version 1.5.3.30 was used to process the raw files and search was conducted against our in house *Saccharomyces cerevisiae* complete/reference proteome database (released from SGD in September 2019), using the Andromeda search engine (Cox *et al*, 2011). For the first search, peptide tolerance was set to 20 ppm while for the main search peptide tolerance was set to 4.5 pm. Isotope mass tolerance was 2 ppm and maximum charge to 7. Digestion mode was set to specific with trypsin allowing maximum of two missed cleavages. Carbamidomethylation of cysteine was set as fixed modification. Oxidation of methionine and phosphorylation of serine, threonine and tyrosine were set as variable modifications. For quantification we set “Reporter Ion MS3” and we chose the 10-plex labels as provided by MQ. We applied the corrections factors provided by the manufacturer.

### Data analysis in R

All data analysis was performed using the proteinGroups.txt and Phospho(STY)Sites.txt files from MaxQuant using R (v 4.2.3) within the RStudio environment. The R scripts will be made available on github. Common yeast gene names were added to the tables by matching the systematic OLN to the common gene names found in the table from https://ftp.uniprot.org/pub/databases/uniprot/knowledgebase/complete/docs/yeast.txt For proteins, the “Reporter.intensity.corrected.” columns in the proteinGroups table corresponding to the measurements from the non-phospho-enriched (“N”) samples were analysed. For phospho-sites, the “Reporter.intensity.corrected.” columns in the Phospho(STY)Sites table corresponding to the measurements from the phospho-enriched (“PE”) samples were analysed. Only singly phosphorylated site intensities (ie Reporter.intensity.corrected columns with suffix “___ 1”) were analysed.

### Analysis of 4 TMT10 prophase block-release timecourses

Proteins or sites marked as contaminants or reverse-matching by MQ were filtered out. Missing values (zeros) were changed to NAs so they were not considered during normalization and batch correction. After these procedures, missing values were converted back to true zeros.To normalise between TMT reporter channels, all values in each reporter column were multiplied by a scaling factor consisting of the median intensity of all reporter columns from all 4 TMT10 timecourse experiments divided by the median reporter intensity of that individual reporter column. For phospho-site normalisation to protein level, the phospho-site intensities from the “PE” reporter columns in the Phospho(STY)Sites table were divided by the corresponding reporter intensity of the same protein from the “N” reporter columns from the proteinGroups table and multiplied by 1000. Normalised protein and phospho-site intensities were then log2 transformed and batch effect correction carried out using the limma package removeBatchEffect function. Following this, log2 transformation was reversed by antilog 2. At this point NA values were changed back to true zeros. All subsequent analysis was carried out using these median-normalised and batch corrected intensities.

To be considered part of the final wild-type or *spo13Δ* timecourse dataset, proteins were required to be detected, having a reporter intensity value >0, in at least one timepoint of both of the two biological replicates of the timecourse. For phospho-sites, the phospho-site was required to have >0 reporter intensity in at least two sequential timepoints in both of the biological replicate timecourses and have a localization probability >0.75. Only phospho-sites that could be normalised to corresponding protein level in all 10 timepoints were analysed further. To be considered part of the final combined wild-type and *spo13Δ* timecourse dataset, proteins and phospho-sites were required to meet thresholds for both individual strain datasets.

To be called significantly dynamic from time 0, a protein or phospho-site needed to meet a fold change and t-test p-value cutoff that considered the abundances in both of the two TMT10 biological replicates of the timecourse. 9 pairwise comparisons between time 0 and each subsequent timepoint were considered. The protein or phospho-site needed to meet the following conditions: First, for at least 1 of the 9 comparisons, it must have a median fold change of greater than 1.5 or less than 1/(1.5). Second, for the same timepoint comparison that meets the fold change cutoff, the difference in abundance needed to be considered significant and consistent between the replicates by returning t-test p-value of < 0.05. The t.test() function from the stats package was used with default parameters, running a Welch’s t-test.

To be considered significantly different abundance between strains, protein or phospho-site abundances at matched timepoints were compared (ie t45 abundance in wild type vs t45 abundance in *spo13Δ*) and the same fold change >1.5 or <1/(1.5) and t-test p-value <0.05 thresholds were applied.

Proteins or phospho-sites which had 0 intensity at one timepoint and >0 intensity at the other timepoint to be compared were allowed to meet the fold change cutoff for being called significantly dynamic from time zero or significantly different between strains. 7% of the wild type proteins and 4% of the combined wild-type and *spo13Δ* protein dataset were considered significantly dynamic from time zero based on a 0 vs >0 comparison and a t-test p-value < 0.05. 29% of wild-type phospho-sites and 17% of the combined wild-type and *spo13Δ* phospho-sites were considered dynamic from time zero based on a 0 vs >0 comparison and a t-test p-value <0.05. 16% of proteins and 23% of phospho-sites were called significantly different between wild type and *spo13Δ* based on 0 vs >0 comparisons as well as a t-test p-value < 0.05. Fold changes from 0 vs >0 comparisons, which are Inf, were imputed by replacing the zeros with the minimum value in the corresponding total dataset and re-calculating the median fold changes. The median-normalised, batch-corrected intensities for each of the two biological replicates of each strain timecourse were averaged together for downstream analyses. This means that the intensities of proteins or phospho-sites which had zero intensity at one timepoint in one replicate were reduced by half.

In the metaphase to anaphase analysis heatmaps, fold changes outside the bounds of the central colour scale were “trimmed” to facilitate seeing the trend in the central data; Fold changes >2.2 were trimmed to 2.2 and fold changes <0.2 were trimmed to 0.2. In the metaphase to anaphase GO term analysis barplots, if a protein increased after the metaphase timepoint, the p-value was plotted as +log10(p-value). However, if the protein decreased after metaphase, the -log10(p-value) was plotted.

For the metaphase to anaphase protein and phospho-site analysis, metaphase I was defined as 75 min and metaphase II as 120 min. Because 135 min was the peak of anaphase II according to the spindle immunofluorescence and phospho-Net1 profile (Figure 1A and 1E), the timepoint just before that, 120 min, was considered metaphase II. Proteins or phospho-sites were called significantly changing from metaphase if they reached >1.5 or < 1/(1.5) fold change from the metaphase timepoint and were significantly different abundance from the metaphase timepoint by t-test p-value < 0.05.

For the metaphase to anaphase phospho-site analysis, metaphase I scaled values for each phospho-site were calculated by dividing all 10 timepoint intensities by the intensity at the 75 min timepoint. Metaphase II scaled intensities for each phospho-site were calculated by dividing all 10 timepoint intensities by the intensity at the 120 min timepoint. In the lineplots, the median and 25-75% quartile range of the metaphase scaled intensities of motif-matching sites are plotted.

For all other lineplots, “scaled intensity” means scaled to the mean. To be mean-scaled, it means that each unique protein or phospho-site was divided by its own mean intensity of all 10 timepoint values, including zeros. For the wild-type and *spo13Δ* combined dataset line plots, protein or phospho-sites were scaled to the mean of both the wild type and *spo13Δ* timecourses together, allowing for inter-strain differences in abundance to be preserved. For summary lineplots (referring to more than one protein or phospho-site), the median and 25-75% quartile range of the mean-scaled intensities are plotted.

### Metaphase I arrest TMT10 experiment analysis

Proteins or sites marked as contaminants or reverse-matching by MQ were filtered out. Missing values (zeros) were changed to NAs so they were not considered during normalization. After normalization, missing values were converted back to true zeros. To normalise between TMT reporter channels, all values in each reporter column were multiplied by a scaling factor consisting of the median intensity of all 10 reporter columns divided by the median reporter intensity of that individual column. For phospho-site normalisation to protein level, the phospho-site intensities from the “PE” reporter columns in the Phospho(STY)Sites table were divided by the corresponding reporter intensity of the same protein from the “N” reporter columns from the proteinGroups table and multiplied by 1000. Only proteins with >0 normalised reporter intensity in at least 3 replicates of all 3 strains were considered part of the final dataset. Only phospho-sites with >0 normalised reporter intensity in at least 3 replicates of all 3 strains and which had a localisation probability >0.75 were considered part of the final dataset. Significant differences in abundance between proteins and phospho-sites between strains were called according to a fold change and t-test p-value cutoff similar to the timecourse experiments. The protein or phospho-site needed to vary between the two strains by a median fold change >1.5 or <1/(1.5), considering all 3 or 4 replicates of each strain. Additionally, the difference in abundance between the two strains, considering all replicates, needed to return a t-test p-value <0.05. The base R t.test() function was used with default parameters, running a Welch’s t-test.

### Hierarchical clustering

Hierarchical clustering was performed using hclust function and cutree functions from the stats package along with the pheatmap package for visualization. Ward’s method was used in the hclust function (method=”ward.D2”). The number of clusters k was determined by identifying the inflection point in a plot of within groups sum of squares vs the number of clusters (ie the ‘elbow method’).

### GO term enrichment analysis

GO term enrichment analysis was performed using the gprofiler2 package with organism = ‘scerevisiae’ and otherwise default settings. This runs a hypergeometric test followed by correction for multiple testing. GO terms were considered significantly enriched if the p-value < 0.05.

### Phospho-site motif logo analysis

The ggseqlogo package was used to make standard sequence logos. The IceLogo software (Colaert et al 2009) was used to make IceLogos using a custom background list of sequences. The percent difference between the two sequences was plotted, with the p-value cutoff set to 0.05. Kinase consensus motifs were all based on those reported in (Mok *et al*, 2010), with the exception of the motif for Mec1^ATR^ defined in (Kim *et al*, 1999) and the motif for Ime2 reported in (Holt *et al*, 2007).

### Fisher tests

Fisher tests were run via the fisher.test() function from the stats package. P-values < 0.001 and were considered significant with three asterisks, p-values < 0.01 were considered significant with two asterisks, and p-values < 0.05 were considered significant with one asterisks.

For the Fisher tests on clustering results in Figure 4 and Figure 9, the number of motif matching sites in selected clusters was compared with the number of motif matching sites in all clusters including the selected cluster (total). For the Fisher tests accompanying the IceLogo analyses in Figure 8, the number of motif matching sites in the selected non-overlapping subsets was compared, to match the subset lists used to make the IceLogos.

## Acknowledgements

We gratefully acknowledge the Wellcome Centre for Cell Biology Proteomics Core for mass spectrometry support. We are grateful to Georg Kustatcher for helpful discussions, to Shaun Webb for coding advice, and Lucia Massari, Gerard Pieper and Menglu Wang for comments on the manuscript. We are grateful to Sandra Touati and Katja Wassmann, and to Joao Matos for sharing results prior to publication. This work was funded through a Wellcome Investigator award to ALM [220780], a Wellcome and Royal Society Sir Henry Dale Fellowship and a Wellcome Multi-User Equipment Grant to TL and core funding for the Wellcome Centre for Cell Biology [203149].

## Author Contributions

Conceptualization – LK and AM; Data curation –LK and CS; Funding Acquisition – TL and AM; Formal Analysis – LK and CS; Investigation – LK, CS, VK; Software – LK; Supervision – TL and AM; Visualization – LK and AM; Writing – original draft – LK and AM; Writing – review and editing – LK and AM with input from all authors

---

**Figure EV1.**
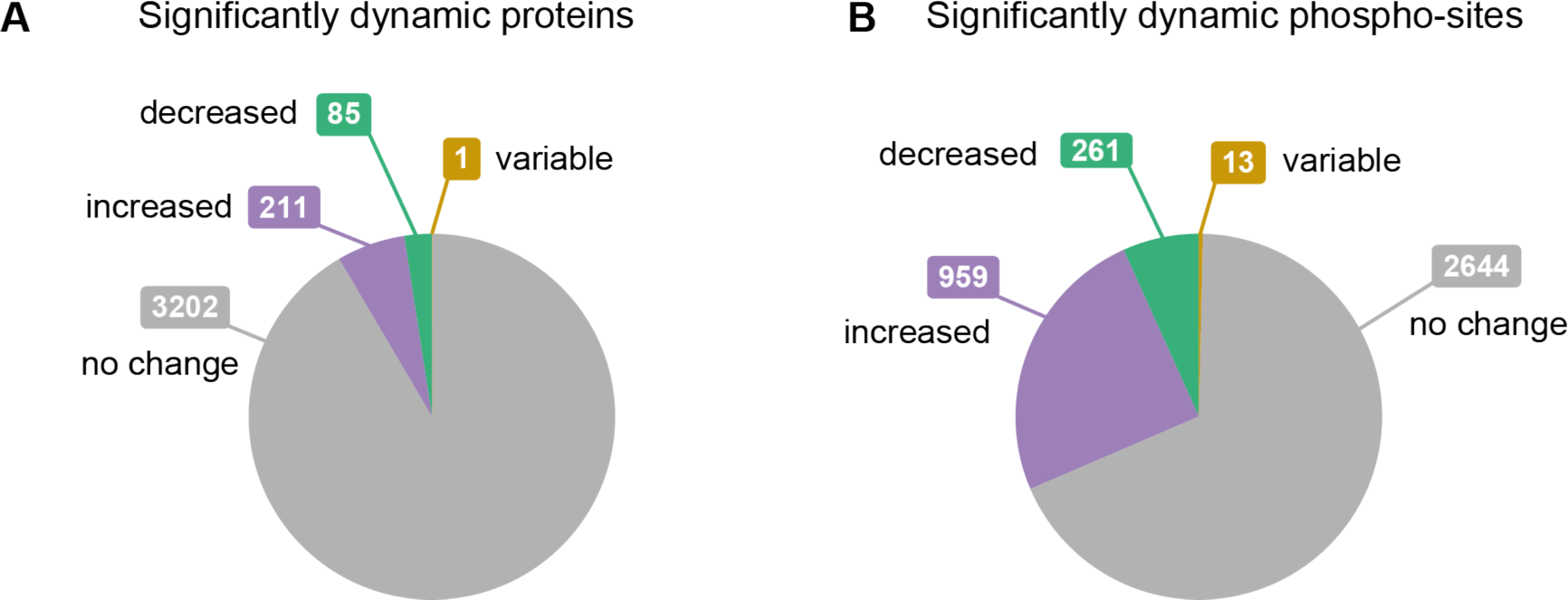
A greater fraction of phospho-sites than proteins are dynamic. A. Proportion of proteins that significantly increased, decreased or had a variable trend from time zero. See Appendix Table S2 for protein identities. B. Proportion of phospho-sites that significantly increased, decreased or had a variable trend from time zero. See Appendix Table S4 for list of phospho-sites in each category.

**Figure EV2.**
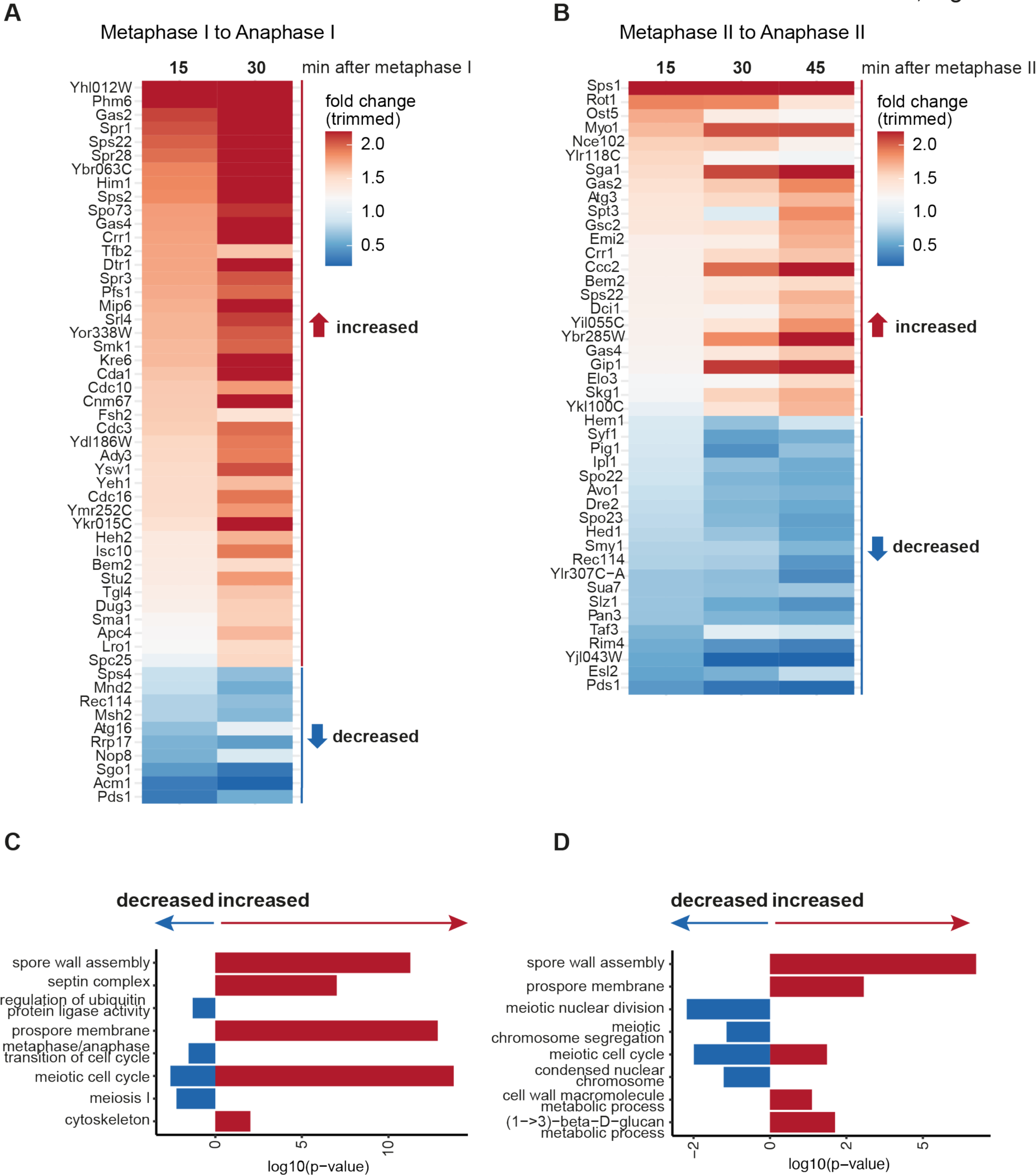
Protein dynamics at the metaphase to anaphase transition in meiosis I and II. A. Fold change in protein abundance of proteins that significantly change from metaphase I to anaphase I. B. Fold change in protein abundance of proteins that significantly change from metaphase II to anaphase II. C. GO term analysis of proteins from A. D. GO term analysis of proteins from B.

**Figure EV3.**
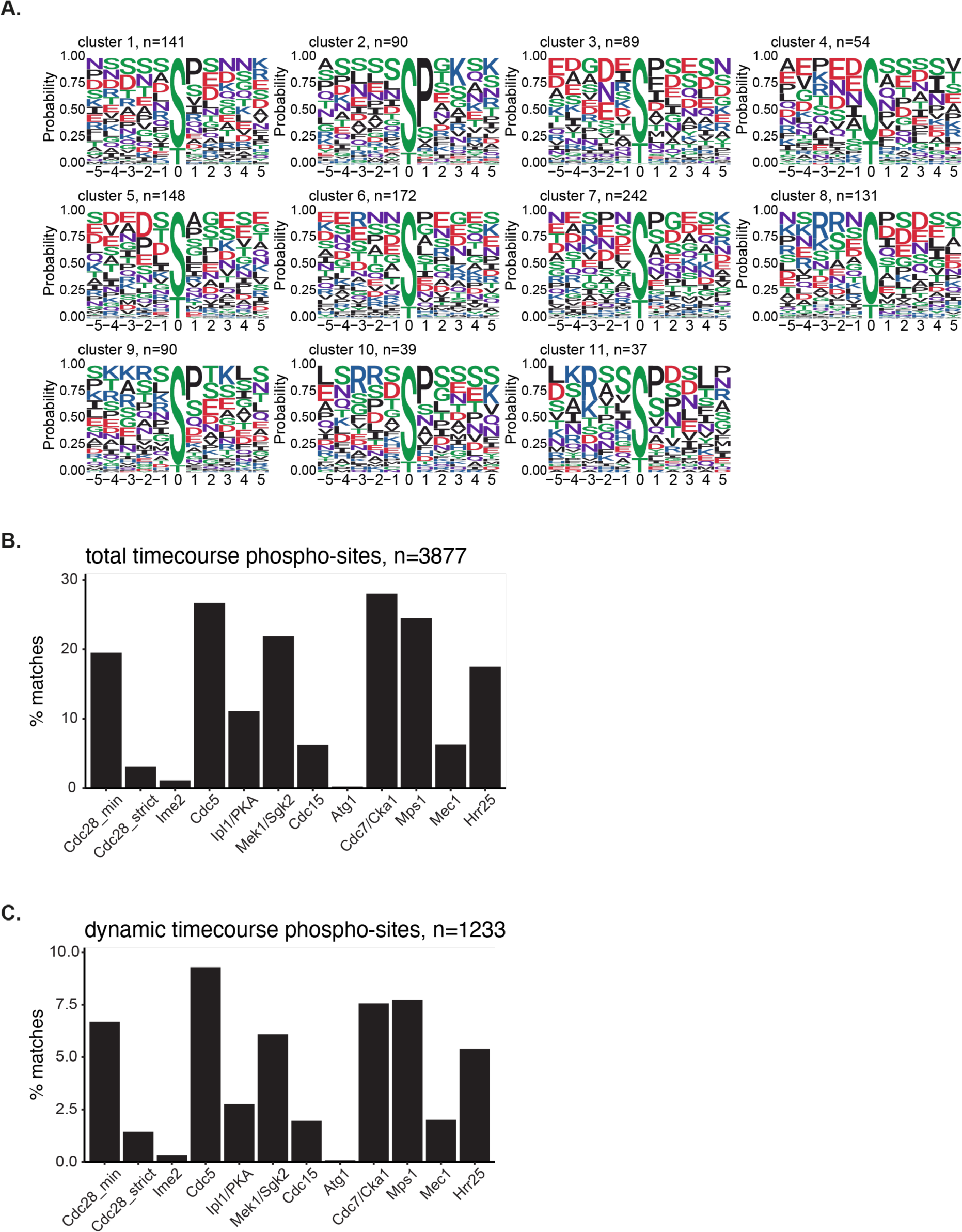
Motif enrichment analysis of phospho-sites across the meiotic divisions timecourse. A. Motif logos of the 11 phospho-site clusters presented in Fig 3 and EV4. B. Bar graph of the percent of motif matching sites for each of the 12 selected kinase motifs in the total timecourse dataset n=3877. C. Bar graph of the percent of motif matching sites for each of the 12 selected kinase motifs in the sites that were dynamic from time zero (including all 11 clusters from Fig 3 and EV4) n=1233.

**Figure EV4.**
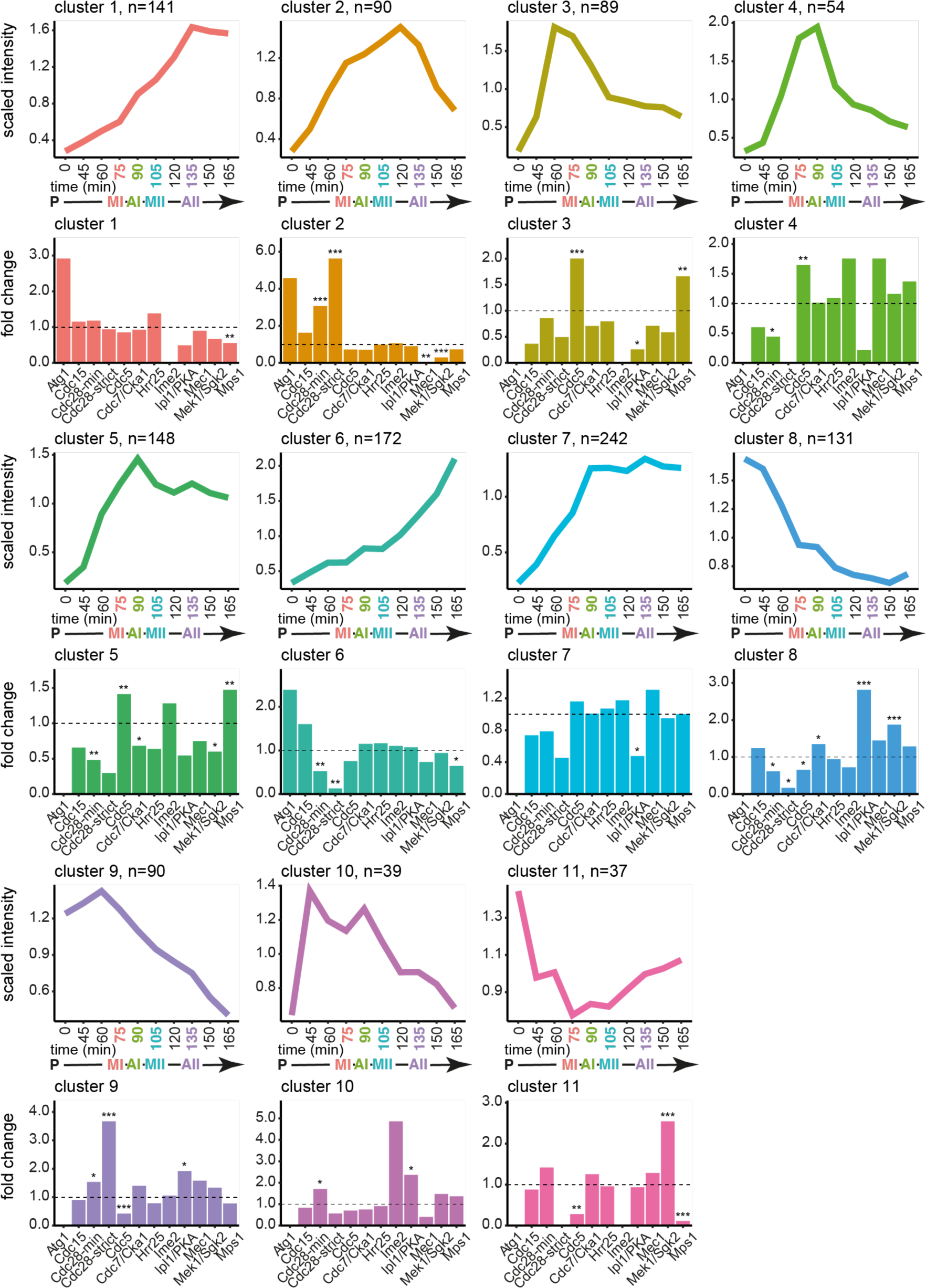
Enrichment of kinase consensus motifs for clusters of dynamic phosphorylation sites. Median lineplots of all clusters from Figure 3A and kinase motif enrichment analysis bar graphs. Asterisks represent p-value from Fisher’s exact test (p<0.001 = ***, p<0.01 = **, p<0.05 = *).

**Figure EV5.**
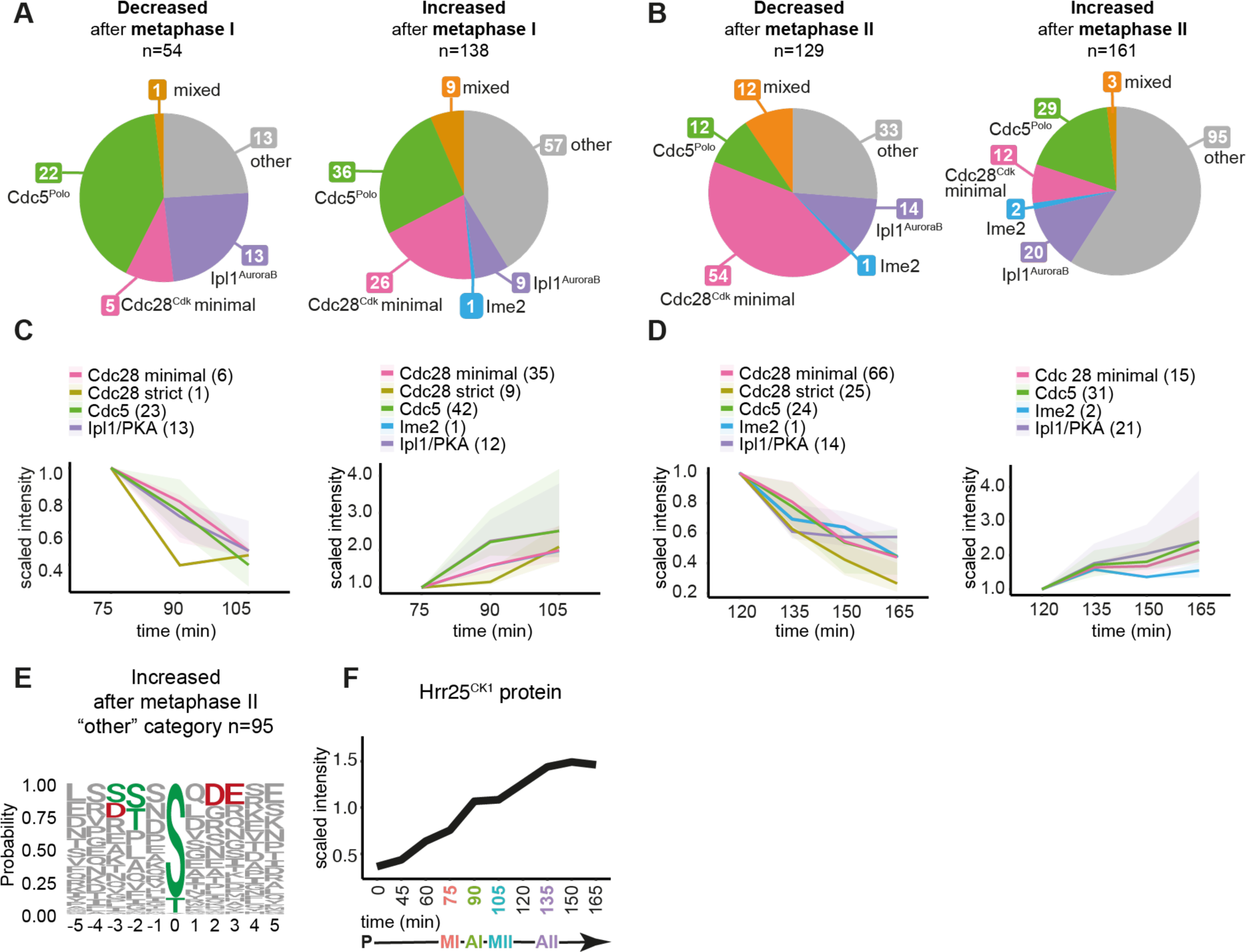
Phosphorylation dynamics at the metaphase to anaphase transitions in meiosis I and II. A. Motifs matching phospho-sites that significantly decrease (left) or increase (right) at the metaphase I to anaphase I transition. B. Motifs matching phospho-sites that significantly decrease (left) or increase (right) at the metaphase II to anaphase II transition. C. Median change of motif matching phospho-sites decreasing (left) or increasing (right) from metaphase I to anaphase I. Abundance scaled to 75min (metaphase I). D. Median change of motif matching phospho-sites decreasing (left) or increasing (right) from metaphase II to anaphase II. Abundance scaled to 120 min (metaphase II). E. Motif logo of phospho-sites that are increased after metaphase II from B (right), which do not match any of the selected motifs, from the “other” category n=95. G. Abundance of Hrr25^CK1^ protein rises in meiosis II.

**Figure EV6.**
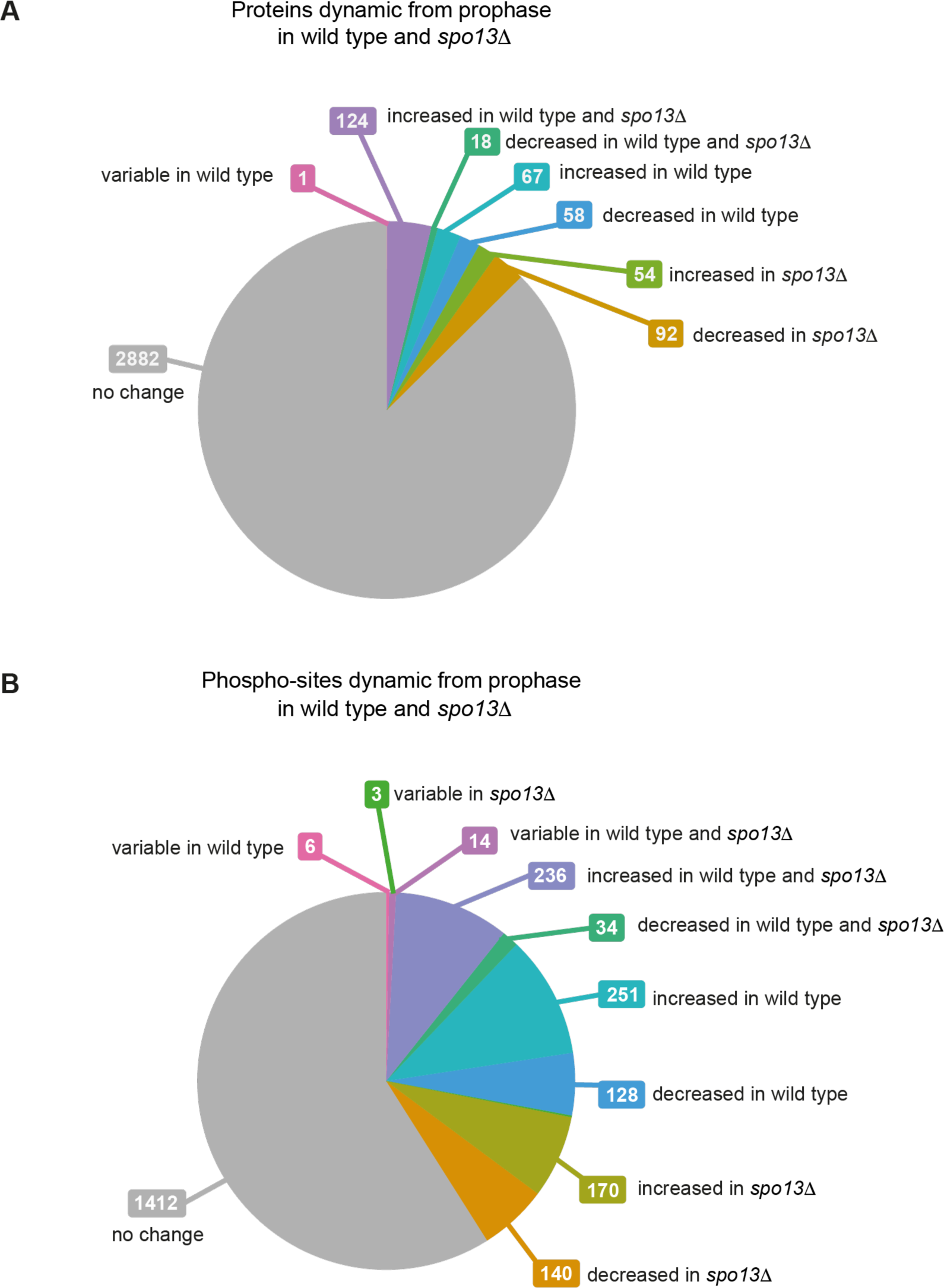
Proportion of dynamic proteins and phospho-sites from prophase in wild type and *spo13Δ*. A. Proportion of proteins that significantly change, increasing or decreasing from time zero, in wild type, *spo13Δ* or both strains. B. Proportion of phospho-sites that significantly change, increasing or decreasing from time zero, in wild type, *spo13Δ* or both.

**Figure EV7.**
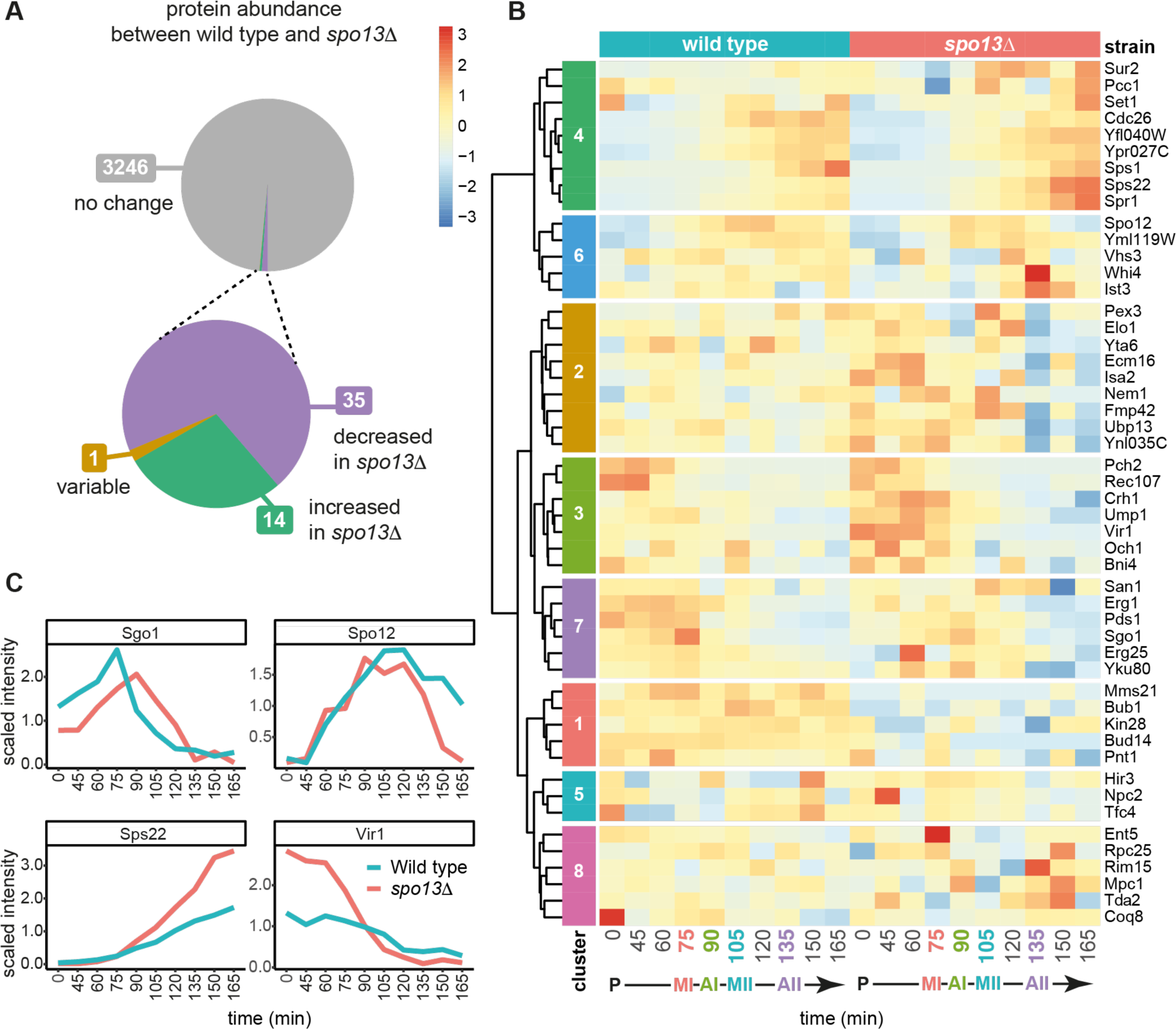
Proteins with significantly different abundance in wild type versus *spo13Δ* cells. A. Proportion of proteins which significantly vary between wild type and *spo13Δ* at matched time points. See also Appendix Table S6. B. Hierarchical clustering of significantly different proteins, n=50, between wild type and *spo13Δ* across the meiotic divisions. See also Appendix Table S7. C. Median abundances of selected proteins that significantly differ between wild type and *spo13Δ* from B.

**Figure EV8.**
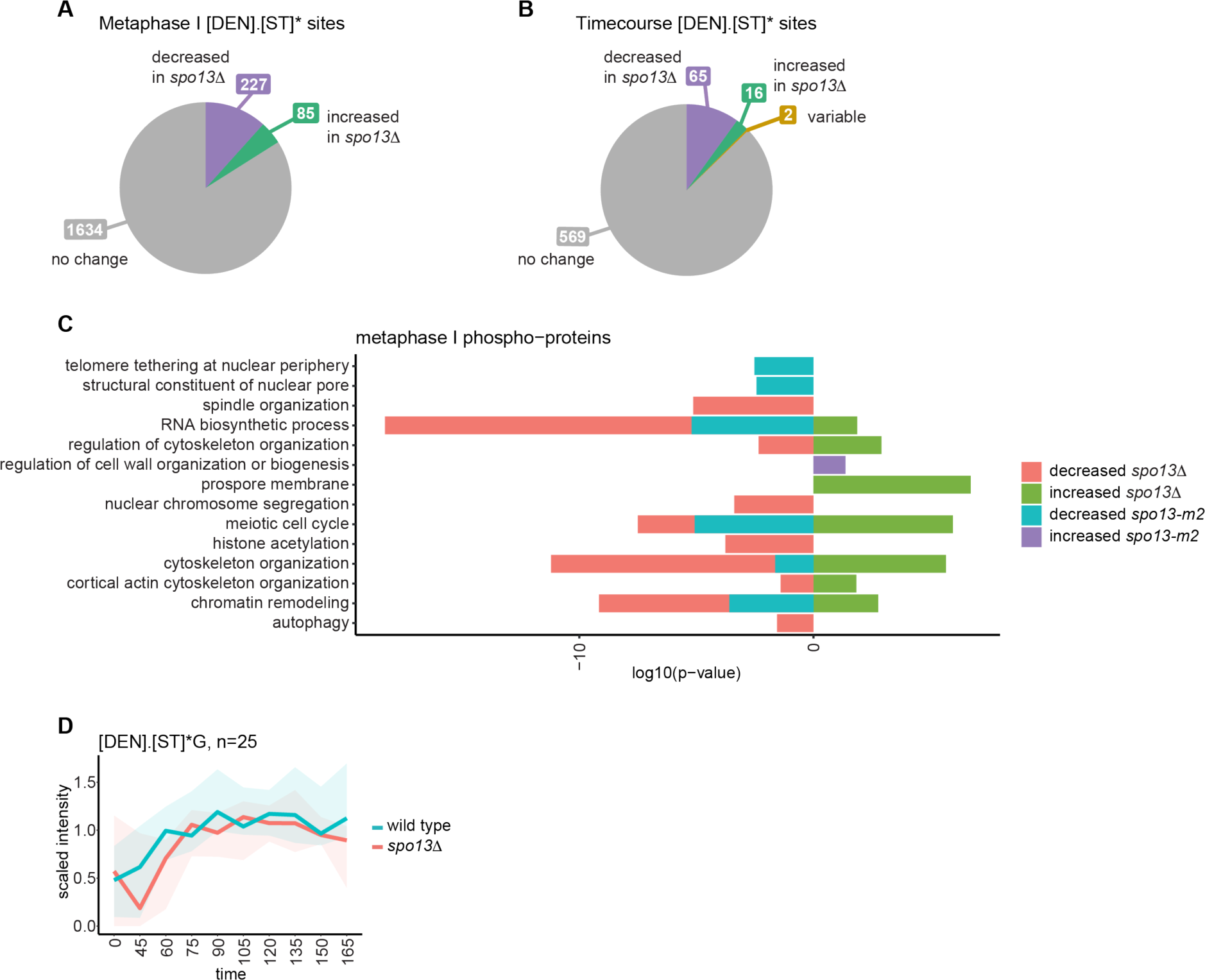
Comparison of Polo kinase motif phosphorylation between wild type and *spo13Δ*. A. Proportion of [DEN]x[ST]* motif matching phospho-sites with significantly different abundance in *spo13Δ* versus wild type in metaphase I arrested cells. B. Proportion of [DEN]x[ST]* motif matching phospho-sites with significantly different abundance in *spo13Δ* versus wild type in the meiotic timecourse experiments. C. GO terms enriched among proteins with significantly different phosphorylation in *spo13Δ* or *spo13-m2* versus wild type. D. Abundance of phospho-sites matching the [DEN]x[ST]*G motif among sites detected in both replicates of wild type and *spo13Δ* across the timecourse.

