## Supplemental data for "Rewiring of the phosphoproteome executes two meiotic divisions"

**Supplementary Information**

**Supplementary Figures**

Koch, Appendix Figure S1

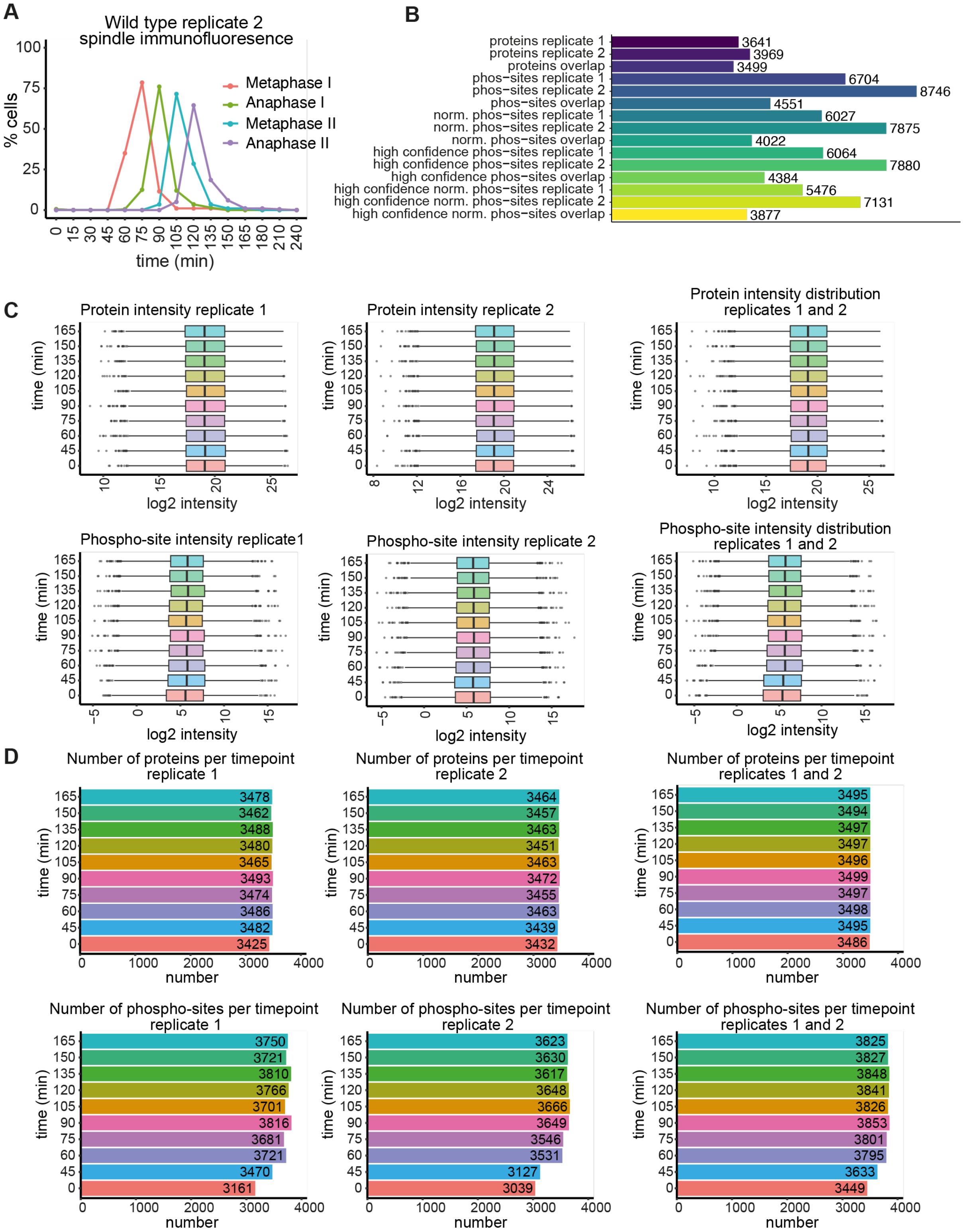


**Appendix Figure S1** A high quality wild type meiotic proteomic and phospho-proteomic timecourse dataset.

A. Morphology of the spindle in the wild type replicate 2 meiotic timecourse.
B. Total numbers of unique proteins and phospho-sites detected for the replicate wild type timecourses. “Norm.” indicates normalized, meaning phospho-site abundance is normalized to corresponding protein level. High confidence indicates localization probability > 0.75.
C. Protein and phospho-site intensity distributions for individual replicates (left two columns) and the combined two-experiment dataset (right column).
D. Protein and phospho-site numbers detected at each time-point for individual replicates (left two columns) and the combined two-experiment dataset (right column).

Koch, Appendix Figure S2

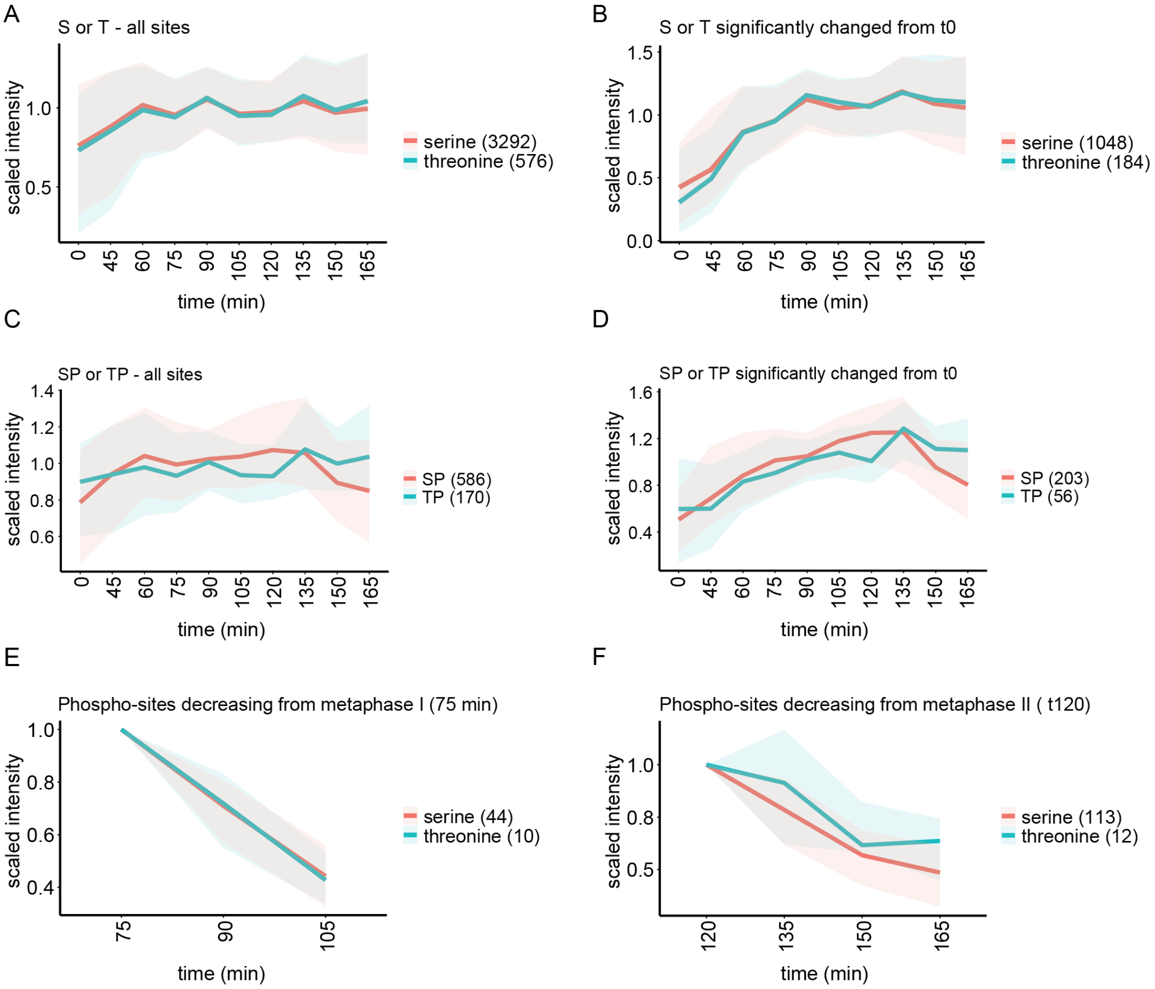


**Appendix Figure S2** Kinetics of the changes in serine and threonine phosphorylation during the meiotic divisions.

A. Scaled median abundance of serine or threonine-directed phosphorylation for all sites in the wild type timecourse.

B. Scaled median abundance of serine or threonine-directed phosphorylation sites that significantly change in abundance from time zero in the wild type timecourse.

C. As for A but including only those phospho-sites that the Cdc28$^{Cdk1}$ minimal consensus [ST]*P.

D. As for B but including only those phospho-sites that match the Cdc28$^{Cdk1}$ minimal consensus [ST]*P.

E. Scaled median abundance for serine or threonine phospho-sites that significantly decrease in abundance after metaphase I (75 min). Scaled to metaphase I abundance.

F. Scaled median abundance for serine or threonine phospho-sites that significantly decrease in abundance after metaphase II (120 min). Scaled to metaphase II abundance.

Koch, Appendix Figure S3

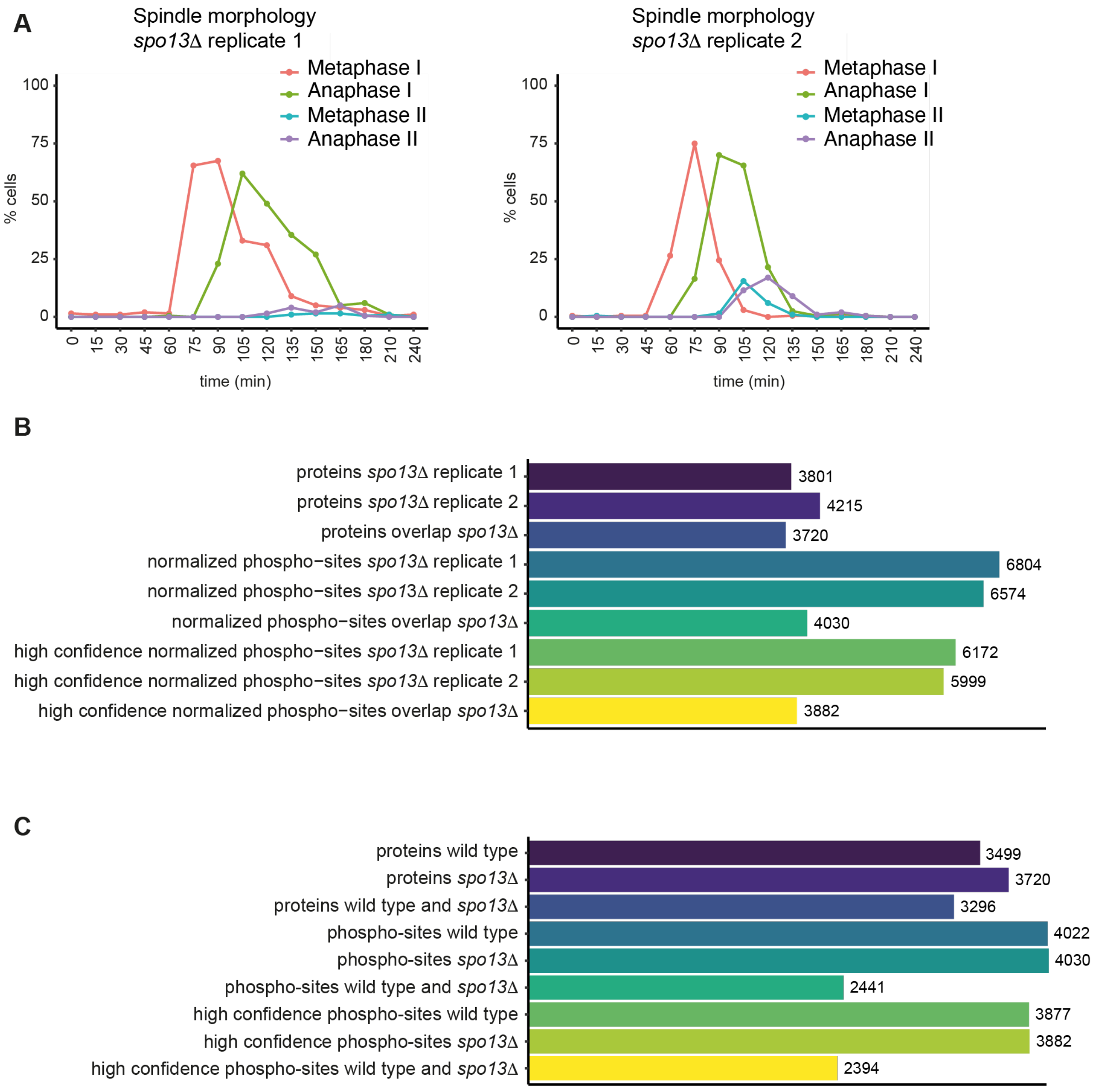


**Appendix Figure S3** The time-resolved phosphoproteome of the *spo13Δ* mutant.

A. Morphology of the spindle in each biological replicate of *spo13Δ* timecourses illustrates the level of synchrony and the single meiotic division of these cells.

B. Total numbers of unique proteins and phospho-sites detected in *spo13Δ* timecourses. Normalized indicates phospho-site abundance is normalized to corresponding protein level. High confidence indicates localization probability > 0.75.

C. Total numbers of unique proteins and phospho-sites detected in the combined wild type and *spo13Δ* dataset. All phospho-site numbers in this plot indicate phospho-sites normalized to corresponding protein level. High confidence indicates localization probability > 0.75.

Koch, Appendix Figure S4

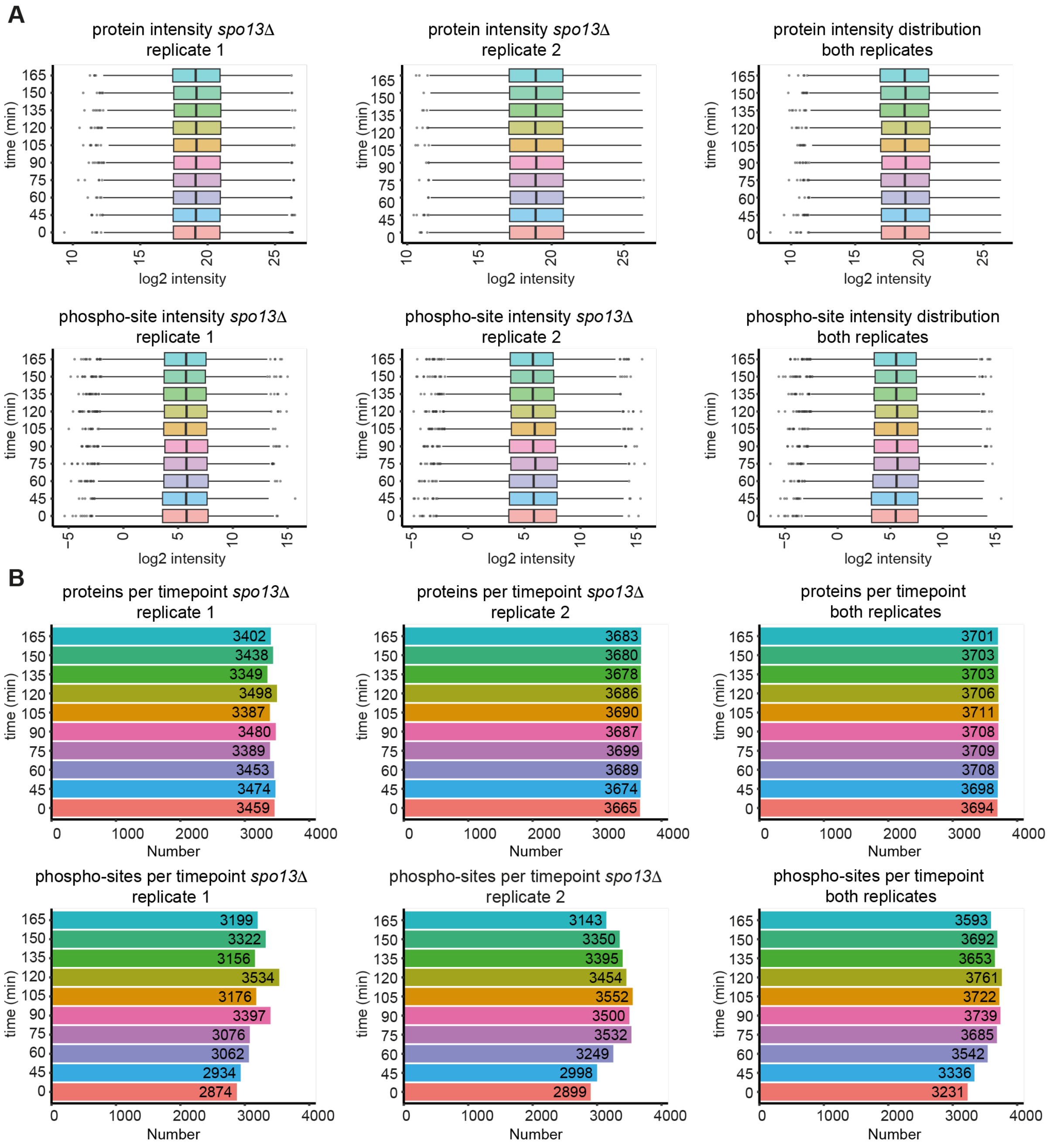


**Appendix Figure S4** Protein and phospho-site intensity distributions and number of proteins detected per timepoint for the *spo13Δ* dataset.

A. Protein and phospho-site intensity distributions for individual replicates (left two columns) and the combined two-experiment dataset (right column).

B. Protein and phospho-site number detected at each timepoint in individual replicates (left two columns) and the combined two-experiment dataset (right column).

Koch, Appendix Figure S5

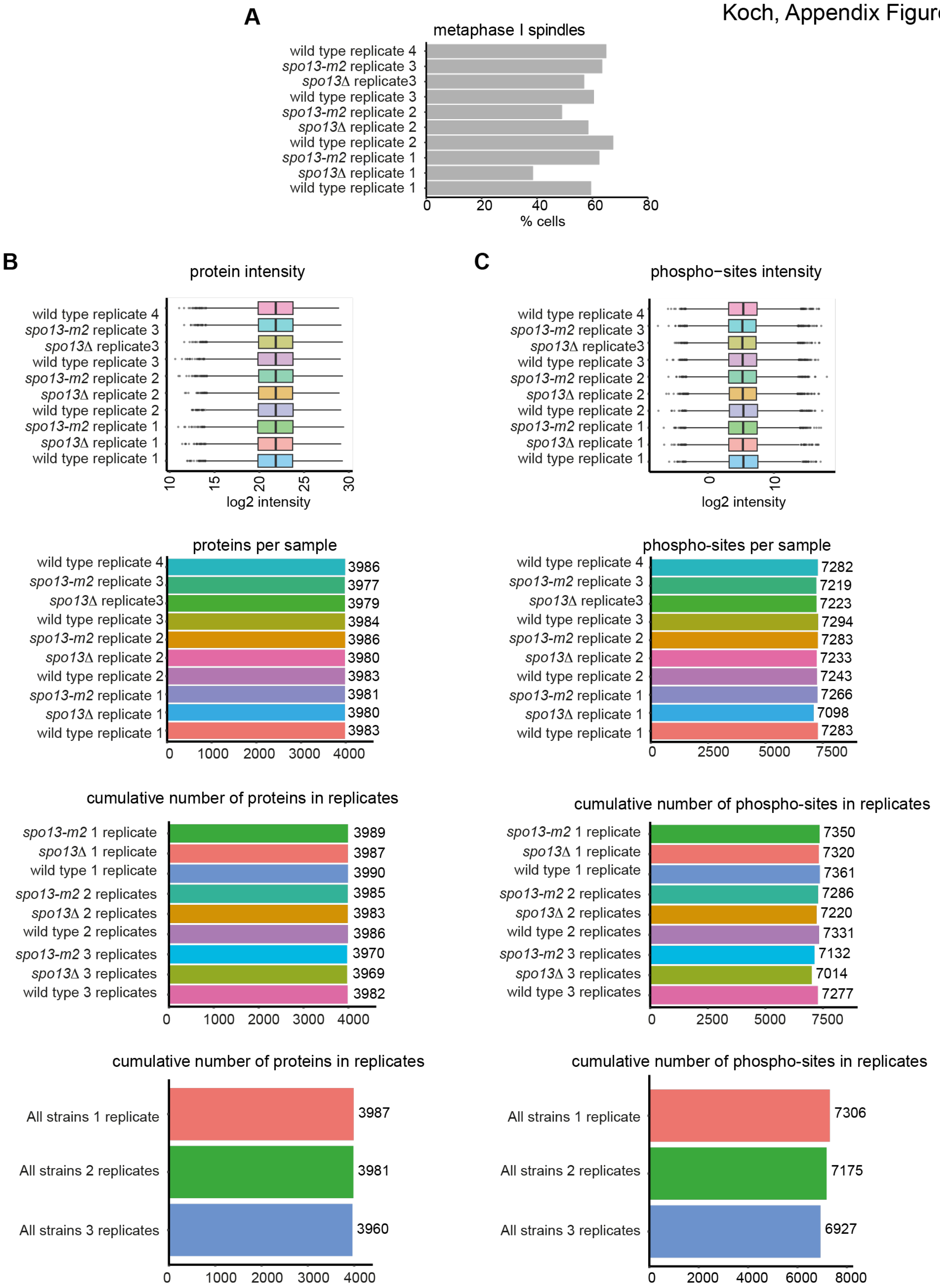


**Appendix Figure S5** Quality control of metaphase I arrest proteomics and phospho-proteomics.

A. Percentage of cells with metaphase I spindle morphology.

B. Protein intensity distribution and number of detections per sample.

C. Phospho-site intensity distribution and number of detections per sample.

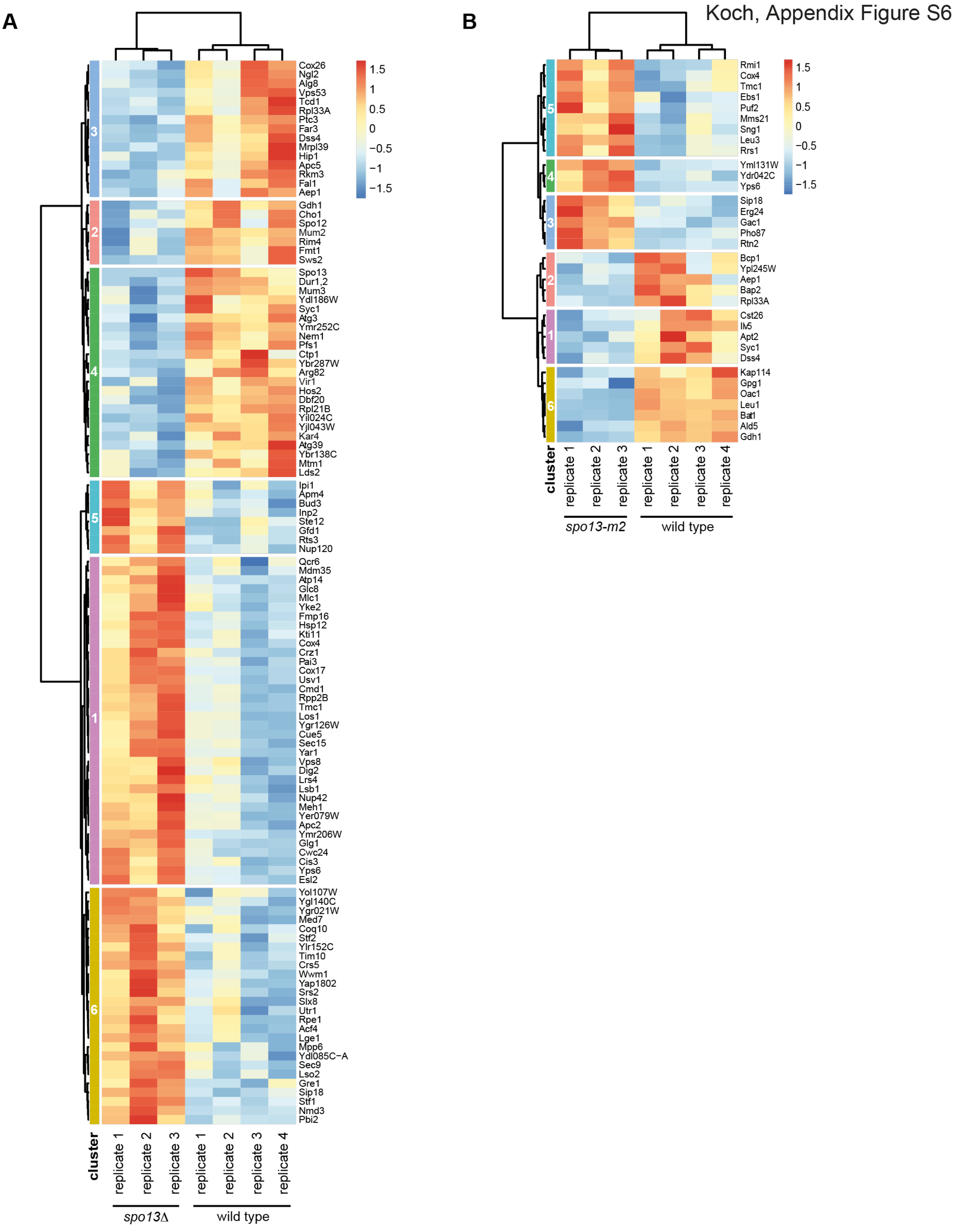


**Appendix Figure S6** Clustering analysis of proteins with significantly different abundance between *spo13Δ* and *spo13-m2* and wild type in metaphase I arrested cells

A. Clustering of proteins with significantly different abundance between wild type and *spo13*Δ in metaphase I arrested cells.

B. Clustering of proteins with significantly different abundance between wild type and *spo13-m2* in metaphase I arrested cells.

**List of Supplementary Tables and Source Data**

**Extended View Table EV1.xlsx**

Kinase consensus motifs used for analysis in Fig 3D, EV3 and EV4.

**Appendix Table S1.xlsx**

Strain Table. Genotype of SK1 budding yeast strains used in this study.

**Appendix Table S2.xlsx**

Significantly dynamic proteins in the wild type timecourse. Lists of proteins in each significance category from Fig EV1A.

**Appendix Table S3.xlsx**

Wild type timecourse protein clustering. Lists of proteins included in each cluster from Fig 2A.

**Appendix Table S4.xlsx**

Significantly dynamic phospho-sites in the wild type timecourse. Lists of phospho-sites in each significance category from Fig EV1B.

**Appendix Table S5.xlsx**

Wild type timecourse phospho-site clustering. Lists of phospho-sites included in each cluster in Fig3A.

**Appendix Table S6.xlsx**

Proteins in each significance category comparing the wild type and *spo13*Δ timecourses from Fig EV7.

**Appendix Table S7.xlsx**

Clustering of proteins significantly different between wild type and *spo13*Δ timecourses from Fig EV7.

**Appendix Table S8.xlsx**

Phospho-sites in each significance category comparing the wild type and *spo13*Δ timecourses from Fig 4.

**Appendix Table S9.xlsx**

Clustering of phospho-sites significantly different between wild type and *spo13*Δ timecourses from Fig 5A-E.

**Appendix Table S10.xlsx**

Proteins significantly different between wild type and *spo13*Δ in metaphase I arrested cells from Fig 6.

**Appendix Table S11.xlsx**

Proteins significantly different between wild type and *spo13-m2* in metaphase I arrested cells from Fig 6.

**Appendix Table S12.xlsx**

Phospho-sites significantly different between wild type and *spo13*Δ in metaphase I arrested cells from Fig 6.

**Appendix Table S13.xlsx**

Phospho-sites significantly different between wild type and *spo13-m2* in metaphase I arrested cells from Fig 6.

**AppendixSourceDataS1.zip**

MaxQuant txt files used for analysis in R.

Timecourse_proteinGroups.txt
Timecourse_phospho (STY)Sites.txt
Metaphase_proteinGroups.txt
Metaphase_phospho (STY)Sites.txt
